# High rates of genome rearrangements and pathogenicity of *Shigella* spp

**DOI:** 10.1101/2020.06.12.147751

**Authors:** Zaira Seferbekova, Alexey Zabelkin, Yulia Yakovleva, Robert Afasizhev, Natalia O. Dranenko, Nikita Alexeev, Mikhail S. Gelfand, Olga O. Bochkareva

## Abstract

*Shigella* are pathogens originating within the *Escherichia* lineage but frequently classified as a separate genus. *Shigella* genomes contain numerous insertion sequences (ISs) that lead to pseudogenization of affected genes and an increase of non-homologous recombination. Here, we study 414 genomes of *E. coli* and *Shigella* strains to assess the contribution of genomic rearrangements to *Shigella* evolution. We found that *Shigella* experienced exceptionally high rates of intragenomic rearrangements and had a decreased rate of homologous recombination compared to pathogenic and non-pathogenic *E. coli*. The high rearrangement rate resulted in independent disruption of syntenic regions and parallel rearrangements in different *Shigella* lineages. Specifically, we identified two types of chromosomally encoded E3 ubiquitin-protein ligases acquired independently by all *Shigella* strains that also showed a high level of sequence conservation in the promoter and further in the 5’ intergenic region. In the only available enteroinvasive *E. coli* (EIEC) strain, which is a pathogenic *E. coli* with a phenotype intermediate between *Shigella* and non-pathogenic *E. coli*, we found a rate of genome rearrangements comparable to those in other *E. coli* and no functional copies of the two *Shigella*-specific E3 ubiquitin ligases. These data indicate that accumulation of ISs influenced many aspects of genome evolution and played an important role in the evolution of intracellular pathogens. Our research demonstrates the power of comparative genomics-based on synteny block composition and an important role of non-coding regions in the evolution of genomic islands.

**Importance:** Pathogenic *Escherichia coli* strains frequently cause infections in humans. Many *E. coli* exist in nature and their ability to cause disease is fueled by their ability to incorporate novel genetic information by extensive horizontal gene transfer of plasmids and pathogenicity islands. The emergence of antibiotic-resistant *Shigella* spp., which are pathogenic forms of *E. coli*, coupled with the absence of an effective vaccine against them, highlights the importance of the continuing study of these pathogenic bacteria. Our study contributes to the understanding of genomic properties associated with molecular mechanisms underpinning the pathogenic nature of *Shigella*. We characterize the contribution of insertion sequences to the genome evolution of these intracellular pathogens and suggest a role of upstream regions of chromosomal *ipaH* genes in the *Shigella* pathogenesis. The methods of rearrangement analysis developed here are broadly applicable to the analysis of genotype-phenotype correlation in historically recently emerging bacterial pathogens.

## Introduction

*Escherichia coli* is likely the best-studied organism, at least on the molecular level. It is widely used to study fundamental aspects of bacterial genomics and is the subject of extensive research as an important component of the normal gut microbiota of vertebrates, including humans. While most *E. coli* strains are harmless, a non-negligible fraction is pathogenic, causing such diseases as diarrhoea, urinary tract infection, or meningitis (1, 2). Another group of pathogens, *Shigella*, which causes a severe form of bacillary dysentery, evolved from *E. coli* (1, 3). *Shigella* spp. are polyphyletic relative to *E. coli*, but the genus name is maintained in part due to the medical tradition (3–5). Nevertheless, from an evolutionary perspective, *Shigella* is just a set of strains causing a specific disease within the broader *E. coli* phylogenetic group.

*Shigella* strains carry a large plasmid (*pINV*) which is essential for virulence (1, 3, 5–8). They also might be distinguished from *E. coli* by their nonmotility with the associated absence of decarboxylated lysine, and by various biochemical characteristics, such as inability to ferment lactose and mucate (3). One more *E. coli* pathovar, enteroinvasive *E. coli* (EIEC), generally exhibits the same pathogenic and biochemical features as *Shigella*, including invasiveness that provided by the *pINV* (1, 3, 5–8). Such phenotypic similarity may be attributed to adaptation to similar environmental conditions as *Shigella* and EIEC spend most of their lifecycle inside eukaryotic cells, while most *E. coli* strains inhabit extracellular space. Thus, EIEC could represent either a *Shigella* ‘prototype’, which could be a precursor for a typical *Shigella*, or a distinct group of pathogenic *E. coli* that have adapted to an intracellular lifestyle but, unlike *Shigella*, have not lost the ability to live outside eukaryotic cells (5, 6, 8).

Acquisition of the virulence plasmid enabling intracellular lifestyle was likely a key event of *Shigella* evolution that facilitated further adaptation. It may have incorporated a variety of events, such as point mutations, suppression of certain genes, deletion of anti-virulence genes, or acquisition of insertion sequences. On the other hand, the intracellular niche may have provided a more relaxed selective pressure due to abundant resources and a lack of competitors (1, 6) that, in conjunction with lower effective population size, would have decreased the negative selection rate (9) and caused substantial changes in its genome arrangement and composition. *Shigella* genomes feature loss or inactivation of many genes, which has been attributed to the relaxation of selection acting on those genes (1). Gene deletions may also contribute to the specialization of bacteria and enable rapid adaptation to different conditions in the host cell (6, 10–12).

The chromosome and plasmids of *Shigella* species contain many insertion sequences (ISs), small mobile DNA fragments that easily translocate within the genome. An analysis of draft *Shigella* genomes demonstrated convergent loss of metabolic pathways by the integration of diverse ISs and pseudogenization by point mutations, often leading to degradation of multiple genes in the same pathway (13). However, the impact of IS elements on genome rearrangements and chromosome evolution has not been studied due to limitations arising from the use of incompletely assembled genome drafts. Indeed, the same repeated IS elements that increase the chromosome instability yield difficulties for the genomes assembly (14, 15).

In turn, repeats accumulation leads to genome rearrangements and changes in the expression of adjacent genes (6, 10, 16–19) affecting the bacterial phenotype. The type of rearrangement depends on the mutual arrangement of the repetitive elements that have been involved in the recombination event. Recombination between inverted repeats leads to inversions, recombination between direct repeats leads to deletions, and recombination between direct repeats during replication leads to duplication (18). Since large deletions, insertions, and duplications are often under negative selection and are rare, inversions are the main drivers of structural changes in bacterial chromosomes (12). The frequency of rearrangements varies and may correlate with the presence of mobile genetic elements (12, 18).

Here, we provide a comprehensive analysis of the complete genomes of *E. coli* and *Shigella* strains based on the construction of synteny blocks and assess the contribution of insertion sequences to *Shigella* genome evolution. We show that *Shigella* genomes experienced exceptionally high rates of intragenomic rearrangements and a decreased rate of homologous recombination in comparison to other, in particular, pathogenic *E. coli* strains. Then we focus on rates of expansion of different ISs families and the patterns of their integration in genomes. Finally, we describe genome rearrangements that have occurred independently in separate lineages, showing convergent evolution of *Shigella*.

## Methods

### Genomes

We used all complete and annotated genomes of *Shigella* and *E. coli* available in `GenBank` as of April 2019 (11). We constructed a phylogenetic tree for *E. coli* strains only (see Methods below) and excluded clones and closely related *E. coli* strains from further analysis to reduce the size of datasets with minimal loss of diversity (Supplementary Figure S1a). Thus, for

*E. coli* strains with identical names on short branches, we selected a random one and removed all others. In particular, we used only one reference genome for *E. coli* K12, *E. coli* О157:H7, *E. coli* O104:H4, *E. coli* O145:H28, *E. coli* BH100, *E. coli* EcoI, *E. coli* O127:H6, *E. coli* O25b:H4, *E. coli* О55:H7, *E. coli* ST540, *E. coli* ST2747, *E. coli* BL21/DE3, *E. coli* Nissle 1917, *E. coli* clone D i2, *E. coli* MRSN and *E. coli* AR strains (the list of excluded genomes is available on GitHub: https://github.com/zseferbekova/ShigellaProject/1Tree/Data/excluded_strains.csv). In total, we analysed 414 complete genomes, including 35 *Shigella* spp., 41 STEC, 31 ExPEC, 8 APEC, 7 ETEC, 3 EPEC, 3 AIEC, 2 EAEC, and 1 EIEC genome (Supplementary Table S1).

### Phylogenetic tree

For the construction of the strains’ phylogenetic tree, we used 238 universal, single-copy orthologous groups found in all 414 genomes. Orthologous groups were constructed using `ProteinOrtho V5·13` (20) with parameters `cov=67 (at least 67% coverage of both proteins in the BLAST alignment)` and `identity=50` (at least 50% identity in the common segments). Then we constructed a nucleotide multiple sequence alignment of genes in each orthologous group using Mafft (21) in the `linsi` mode. We then used RAxML (22) with the `GTR+Gamma` model and 100 bootstrap replicates to construct a phylogenetic tree based on the concatenated alignment of these genes (Figure 1). Finally, we used the GGRaSP (23) R package to divide all strains in the tree into seven clusters corresponding to the standard phylogroups (24). All trees were visualized using online iTOL (25).

**Figure 1.**
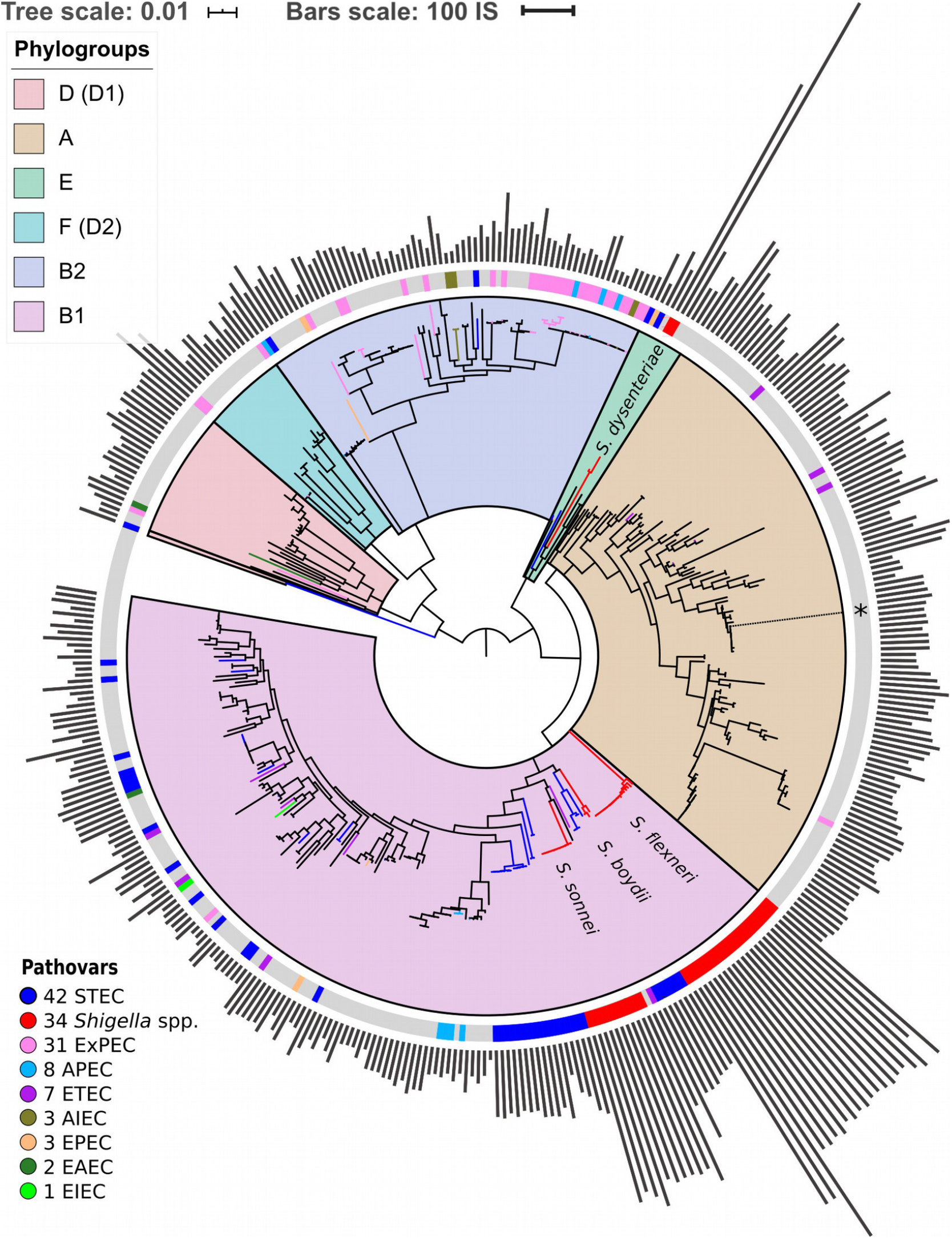
Phylogenetic tree of *E. coli* and *Shigella* spp. The tree is based on the nucleotide alignment of universal single-copy orthologs. Clusters and corresponding phylogroups are shown in different colours; the number of genomes of each pathovar is indicated to the right of the pathovar’s name. Pathogenic *E. coli* strains are marked with coloured bands on the outer circle. The location of the unclassified *Shigella* strain is shown with an asterisk. The bars indicate the number of ISs found by ISsaga in bacterial chromosomes. The tree is unrooted. The tree is also available online (iTOL): https://itol.embl.de/tree/8589127100415001556065531.

Additionally we checked the robustness of the phylogenetic tree with regards to the identity threshold in the definition of orthologs. For that, we reclustered the orthologs with the protein identity threshold `= 95%` and compared the phylogenetic trees (Figure 1, Supplementary Figure S1b). The trees are consistent except for a small number of internal nodes with low bootstrap support. As we do not directly use the fine topology of the phylogenetic tree, that does not affect the results, nor does that influence the chromosome rearrangement analysis, as the latter has been performed directly on nucleotide sequences of the genomes.

### IS elements

Online tool `ISsaga` (26) was used for annotation of IS elements in the chromosome sequences. For general statistical calculations, we used the number of predicted ORFs associated with ISs. For more precise annotation of ISs from different families, we used the number of predicted ISs that could include several ORFs.

### Synteny blocks

The multiple whole-genome alignment for construction of locally-collinear blocks was performed with `SibeliaZ` (27). This new approach was specifically designed to address the performance issues caused by the large number of assembled genomes. It allowed us to scale our analysis to more than 400 genomes while maintaining accuracy. The k-mer size (`-k`) was set to 15, which is recommended by the documentation for bacterial genomes. Next, submodule `maf2synteny` from `Ragout` (28) was used to merge locally collinear blocks into synteny blocks (Supplementary Table S2). This approach is not sensitive to the annotation of genomes and identification of orthologs since it is based on compacted de Bruijn graphs constructed directly for nucleotide genomic sequences. The minimal block size (`-b`) parameter was set to 1000, the simplification parameter (`-s`) was set to `fine` in order to retain the information about small-scale rearrangements. These stringent parameters allowed us to extend the analysis of rearrangements to recent pseudogenes, RNA *gene*s, and conserved intergenic regions. Location of the synteny blocks in the chromosomes was visualised using `Circos` (29).

To infer the number of inversions on the phylogenetic tree we used common single-copy synteny blocks with the block size threshold (`-b`) of 5 kb. In each phylogenetic cluster, we used distance matrices where each element is a number of synteny blocks between given strains. We constructed trees based on the obtained matrices using PHYLIP (30) and the neighbour-joining algorithm.

### Inversions scenario

We reconstructed the history of inversion events using `MGRA` (31). This tool takes as an input a phylogenetic tree and genomes represented as sets of synteny blocks. This analysis included chromosomes of all *Shigella*, related *E. coli* and several representative *E. coli* from each cluster not containing *Shigella* (51 genomes in total).

### Parallel rearrangements

We say that a rearrangement is *consistent* with a tree if we may associate it with a particular branch on a tree, otherwise, we call a rearrangement *parallel*. We test each rearrangement for consistency with a tree with the standard Fitch algorithm (32). This approach allows us to detect the events occurring multiple times in distant clades, in particular, in different *Shigella* lineages.

To analyse inversions, we considered common single-copy blocks. For these 377 blocks, we constructed the breakpoint graph (33, 34) as follows. The graph is built on 377×2 vertices. For each block *B*, we introduce two vertices *B*_H_ and *B*_T_, its head and tail, respectively. If two blocks *B* and *C* are adjacent in genome *g*, the vertices *B*_H_ and *C*_T_ are linked by an edge of colour *g*. We note that since we consider only common blocks and all genomes are circular, the edges of each colour form a perfect matching on the graph vertices. Since some adjacency edges of different colours may link the same pair of vertices, we introduce multi-edges — a multi-edge is a set of parallel edges of different colours. The breakpoint graph for our data contains 754 vertices and 656 multi-edges. If two strains differ from each other by one inversion, this corresponds to a 4-cycle in the breakpoint graph (Supplementary Figure S2ab). For each multi-edge, we split the set of strains into patterns depending on the presence of the corresponding colour edge in the multi-edge (Supplementary Figure S2c). Thus we associated each breakpoint (and each inversion) with a partition of the set of strains into patterns.

To analyse insertions, deletions, and duplications, we considered all blocks which were present in different copy numbers in some strains. For each block, we split the set of strains into patterns so that the copy number of this block in each pattern was the same (i.e. the pattern with strains containing 0 copies of the block, the pattern with strains containing 1 copy of the block, etc). Thus, we associated each copy number variation with a partition of the set of strains into patterns. Then, for blocks whose copy numbers differ in *Shigella* and *E. coli*, we manually classified the evolutionary scenarios based on the occurrence pattern and functional annotation of genes found in the block.

The developed pipeline for the detection of parallel rearrangements is available on GitHub: https://github.com/ctlab/parallel-rearrangements.

### Rates of homologous recombination

For this analysis, we considered genomes from phylogroup B1 as it contains most of the *Shigella* strains and for better resolution used pairwise full-genome alignments constructed using `MAUVE` (35). Then for fragments without gaps, we calculated the number of non-identical columns in each 1 kb segment. Thus, for each pair of genomes, we constructed the distributions of the number of mutations across the genomes.

In the case of strictly vertical inheritance, this distribution would be Poisson with the parameter λ reflecting the time of strain divergence. Fragments transferred horizontally from distant strains would contain more mutations yielding deviation from the Poisson distribution in the form of a heavy tail (36). The latter, being a mixture of the Poisson distributions with unknown parameters may be fitted by the Erlang distribution.

We have used the Python SciPy module (37) to fit all pairwise distributions by the function *F*_λ,*k*,μ,*W*_(*x*) = *W*×**P**_λ_(*x*)+(1–*W*)×**E**_*k*,μ_(*x*), where **P**_λ_(*x*) = *e*^−λ^λ^*x*^/*x*! is the Poisson distribution with parameter λ, **E**_*k*,μ_(*x*) = (*x*/μ)^*k*–1^*e*^−*x*/μ^/(μ(*k*–1)!) is the Erlang distribution with the shape *k* and scale μ (mode=(*k*–1)μ, mean=*k*μ, variance=*k*μ^2^), and the weight *W* in the range [0,1] measures the vertically inherited fraction of genome while (1–*W*) corresponds to horizontally transferred fraction.

This approach extends the one suggested in (36). It averages over all genome segments, and hence is more robust than the approaches based on explicit identification of recombined segments, as the latter are sensitive to uneven evolutionary rates and, moreover, are computationally prohibitive for large-scale analyses. The Poisson parameter λ monotonically increases with the divergence time of the vertically inherited genome fraction, and selecting pairs with the same λ, we obtain a set of strain pairs that have diverged at approximately the same time.

## Results

### Structure of the phylogenetic tree and accumulation of IS elements

We found 238 universal single-copy orthologous groups in 414 genomes (Supplementary Table S1) and used them to construct the unrooted phylogenetic tree (Fig. 1). The structure of the obtained phylogenetic tree recapitulates known *E. coli* phylogroups and supports the hypothesis that *Shigella* spp. included in our analysis evolved several times independently from *E. coli* and are named in accordance with the tree branches (4, 24). One of the *Shigella* genomes (GenBankID: GCA_001596115.1) was unclassified and did not cluster with any described *Shigella* species. Moreover, the source of the sample was lichen, which is highly unusual and unlikely for *Shigella*. Thus, we assumed that in this case the taxonomic annotation was wrong and did not consider this genome as *Shigella*. The only complete and annotated EIEC strain did not cluster with any *Shigella* (38). Other pathogenic *E. coli* strains also did not form any monophyletic clusters.

*Shigella* genomes generally encode more IS elements than non-pathogenic *E. coli* (10). To estimate the density of IS elements, we calculated their number in the chromosomes of all 414 strains (Fig. 1). The number of IS elements in *Shigella* genomes was significantly higher than in the genomes of pathogenic and non-pathogenic *E. coli* strains (Fig. 2a; the Wilcoxon-Mann-Whitney test, *p*-value = 2.4×10^−21^ and 7.8×10^−18^, respectively). Interestingly, the two most sequenced pathovars, STEC and ExPEC, had, respectively, significantly higher (*p* = 1.9×10^−4^) and lower (*p* = 3.5×10^−4^) number of IS elements than the average in non-pathogenic *E. coli* (Fig. 2b).

**Figure 2.**
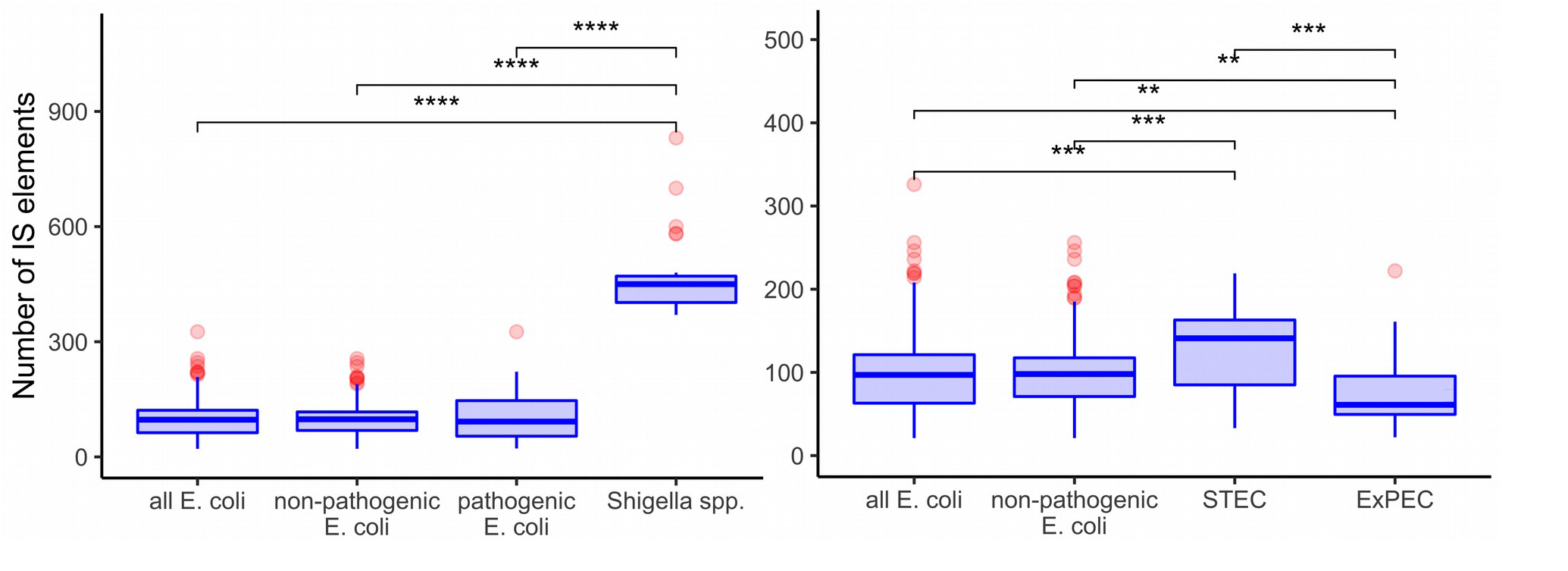

**Comparison of IS numbers** in **(a)** *Shigella* spp., non-pathogenic *E. coli*, other pathogenic *E. coli*, and all *E. coli* (excluding *Shigella* spp); **(b)** two most abundant pathovars — ExPEC and STEC, non-pathogenic *E. coli*, and all *E. coli* (excluding *Shigella* spp). The Wilcoxon-Mann-Whitney test with the Bonferroni correction: (*) *p* ≤ 0.05, (**) *p* ≤ 0.01; (***) *p* ≤ 0.001; (****) *p* ≤ 0.0001.

To test whether the distribution of IS families differed in two clusters of *Shigella* we merged IS elements into larger families and mapped the results on the phylogenetic tree (Fig. 3, 4). We found four IS families (IS1, IS3, IS4, IS91) which were enriched in all *Shigella* in comparison to *E. coli*. Moreover, we detected IS families that were specific for some *Shigella* lineages (IS21, IS110, IS630, IS66). However, as our dataset included only three *S. boydii* and two *S. dysenteriae* genomes, the results for these lineages should be considered as preliminary. By contrast, the *S. sonnei* group was represented by 11 strains and showed a significantly higher number of IS21, IS110 and IS630 elements (the Wilcoxon-Mann-Whitney test, *p* = 1.3×10^−8^, 5.1×10^−7^, 1.9×10^−9^, respectively).

**Figure 3.**
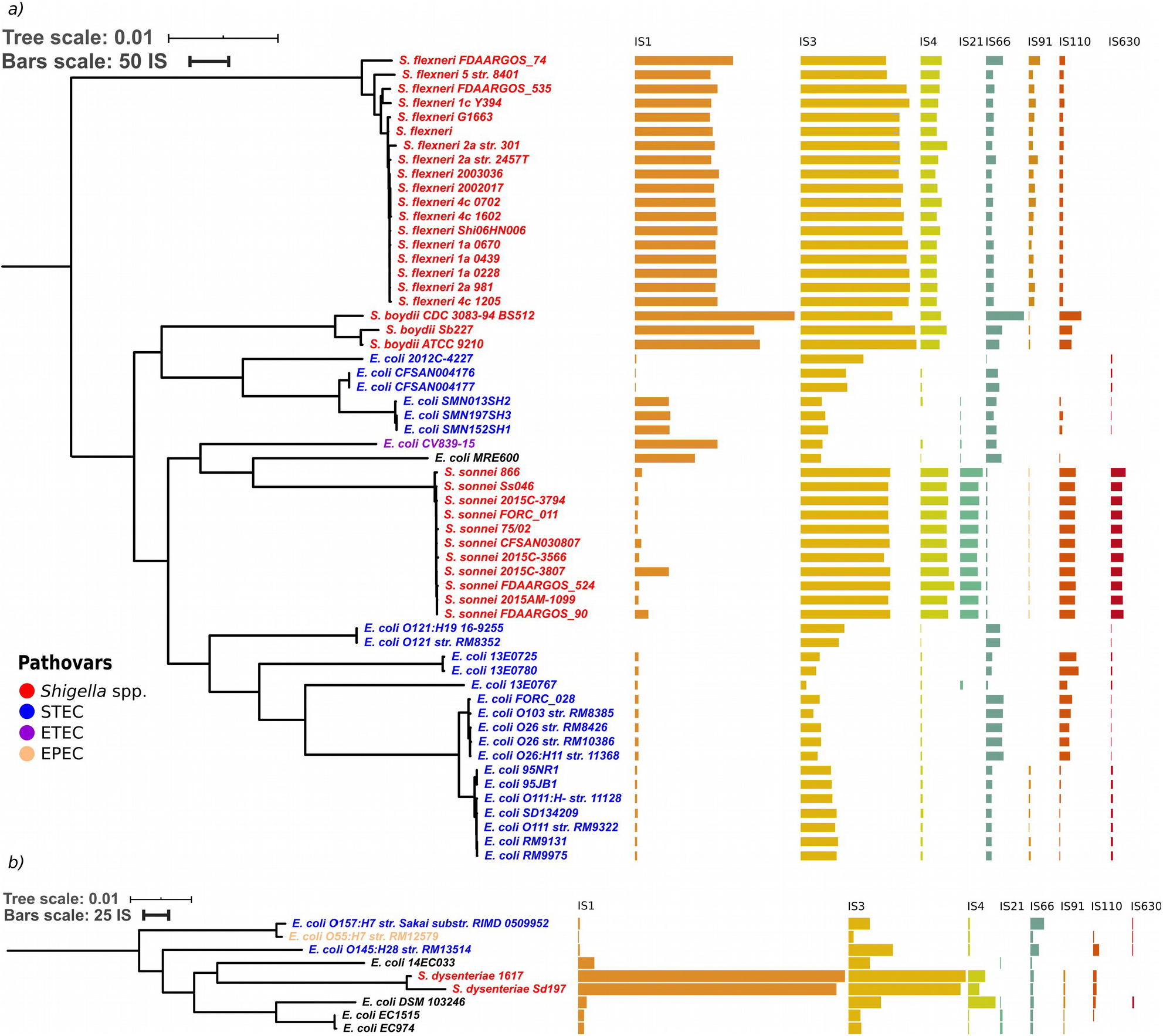

**Distribution of IS families** in **(a)** a part of phylogroup B1 and **(b)** phylogroup E. The bars indicate the number of ISs in each family. The trees are also available online (iTOL): a) https://itol.embl.de/tree/8589127100479001583084903#, b) https://itol.embl.de/tree/8589127100208261560504457#.

**Figure 4.**
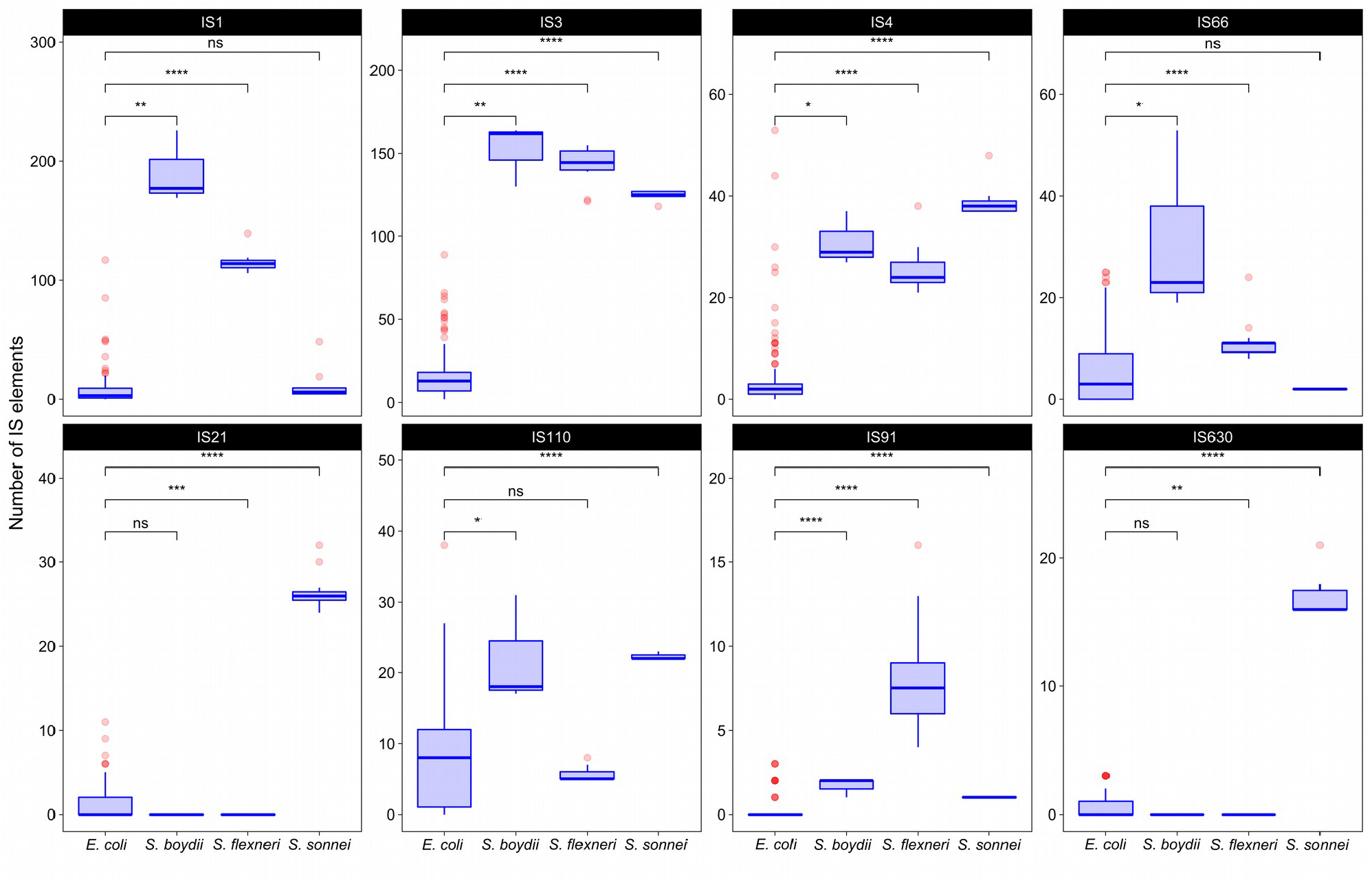
The comparison of selected IS families in *Shigella* and *E. coli* strains from phylogroup B1. The Wilcoxon-Mann-Whitney test with the Bonferroni correction: (*) *p* ≤ 0.05, (**) *p* ≤ 0.01; (***) *p* ≤ 0.001; (****) *p* ≤ 0.0001.

### Composition of synteny blocks

In all studied *Shigella* and *E. coli* genomes we found 4019 synteny blocks. The distribution of the synteny blocks by the number of strains in which they are present has an asymmetric U-shaped form similar to the distribution of gene frequencies in a population, also known as the *U-curve* (Fig. 5a). Only 377 synteny blocks were classified as universal, so that each block was found exactly once in all considered genomes, while other 3642 synteny blocks were not found, or found more than once, in at least one genome. The mean fraction of a chromosome covered by synteny blocks with the length threshold 1 kb was 62%. The universal blocks spanned only 25–29% of the chromosome length and the distribution of these blocks across the chromosomes was not uniform, with long sections not harbouring any universal blocks (Supplementary Figure S3). The comparison of the distributions of common blocks across the chromosomes in different *Shigella* lineages, combined with GC-skew plots, revealed numerous unbalanced genomic rearrangements.

**Figure 5.**
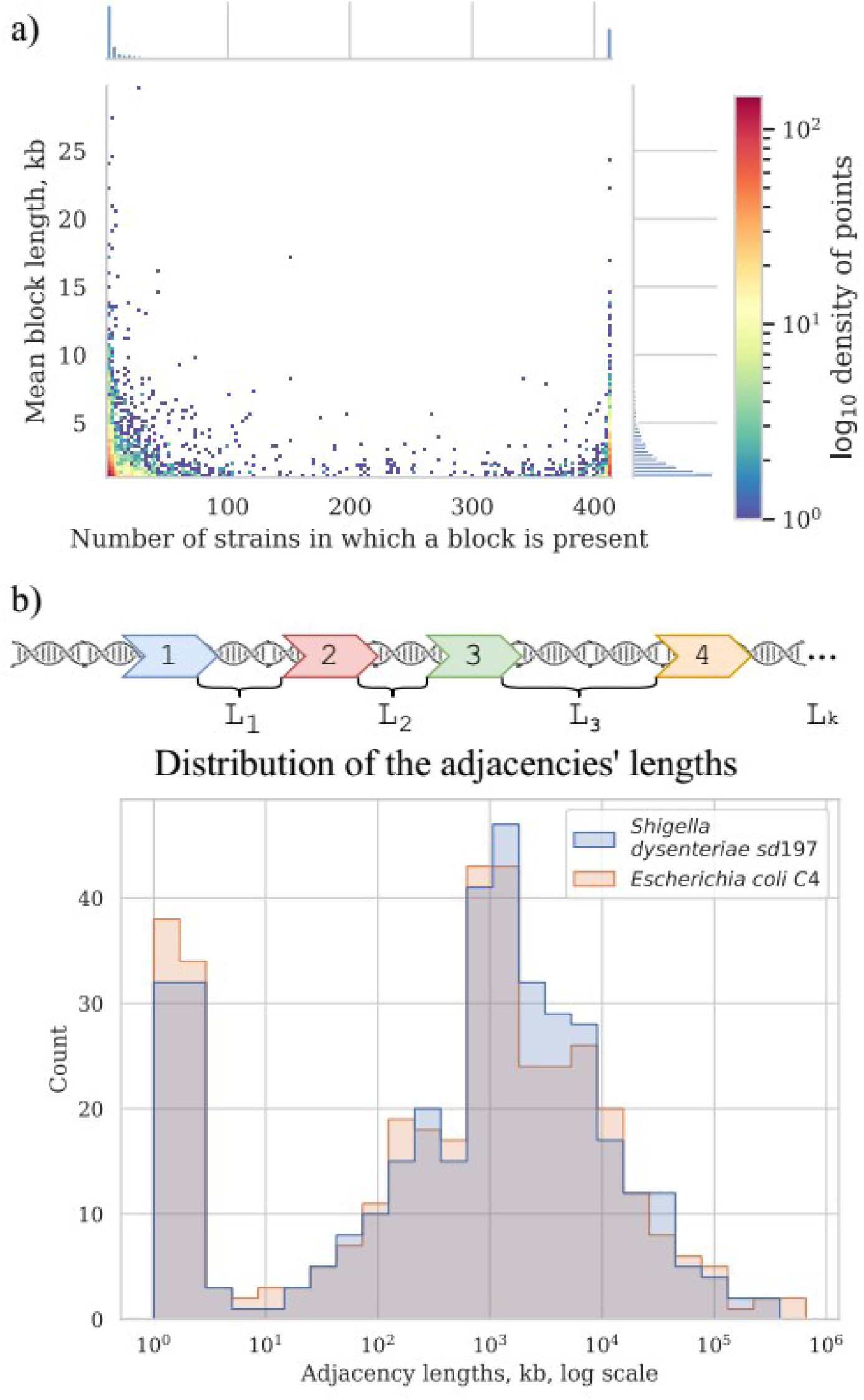
Synteny blocks. **(a)** 2D histogram shows the relationship between the mean block length and the number of strains in which it is present. Color of the point corresponds to the number of synteny blocks with these parameters. *X*-axis histogram shows the frequency distribution of the synteny blocks; *y*-axis histogram shows the length distribution of the synteny blocks **(b)** Distribution of the adjacencies’ lengths in *E. coli* and *Shigella*, in nucleotides. Adjacencies are defined as the chromosome regions between the neighboring universal synteny blocks (see Methods).

Most of the blocks are shorter than 5 kb; some exceptionally long blocks are found among both common blocks and blocks that are present only in few strains (Fig. 5a). Common long blocks are formed by operons of housekeeping genes, the longest ones being (1) 20 genes including DNA polymerase III subunit alpha, components of the complex involved in the assembly of outer membrane proteins and elongation factors; (2) 19 genes including components of the cell division complex and the *mur* operon; (3) 14 genes that are components of the NADH-ubiquinone oxidoreductase complex. The longest rare blocks are formed by recent insertions such as prophages and pathogenicity islands.

The focus on the presence/absence patterns of synteny blocks allowed us to distinguish between genome rearrangements not affecting copy numbers (such as inversions) and those leading to copy number variations (such as insertions, deletions, and duplications). We then separately constructed the breakpoint graph for universal blocks and analysed phyletic patterns of non-universal blocks (see Methods). Thus we identified genetic features that could not be parsimoniously explained by common ancestry.

### Rates of genome rearrangements

*Shigella* genomes are thought to be dynamic due to numerous IS elements that promote non-homologous recombination (6, 16, 17). Thus, the rate of rearrangement in *Shigella* genomes may be higher in proportion to the accumulation of single nucleotide substitutions in comparison to non-invasive pathogenic *E. coli*. To test this possibility we compared the number of syntenic blocks with the number of single nucleotide substitutions in the universal genes for pairs of genomes in each phylogroup. The number of common single-copy synteny blocks between two genomes was inversely proportional to collinearity of the genomes, i.e. more collinear genomes had fewer blocks while more blocks corresponded to genomes with a large number of inversions. While the number of syntenic blocks may not accurately reflect the number of rearrangement events, especially when the number of such events is substantial, our approach provides a lower bound estimate of the number of inversion events. We then plotted the number of syntenic blocks relative to the number of single nucleotide substitutions for each pair of genomes (Supplementary Figures S3, S4).

Indeed, within the same interval of evolutionary distances between strains, in pairs of *Shigella* and *E. coli* strains ratio of a number of syntenic blocks to single nucleotide substitutions was substantially higher, in comparison to pairs of *E. coli* strains (the Wilcoxon-Mann-Whitney test, *p*=2.22×10^−16^) (Fig. 6a). Moreover, for pairs of *Shigella* this ratio is even higher and different in four *Shigella* lineages (Fig. 6b). Thus, genome rearrangements were occurring more frequently in the *Shigella* history compared to *E. coli*. In other pathogenic *E. coli* ratio of a number of syntenic blocks to single nucleotide substitutions did not differ from the average in non-pathogenic strains (Supplementary Figure S4a-d). The unclassified *Shigella* strain from lichen did not differ from *E. coli* strains (Supplementary Figure S4a), further supporting our assumption that this strain has been misclassified.

**Figure 6.**
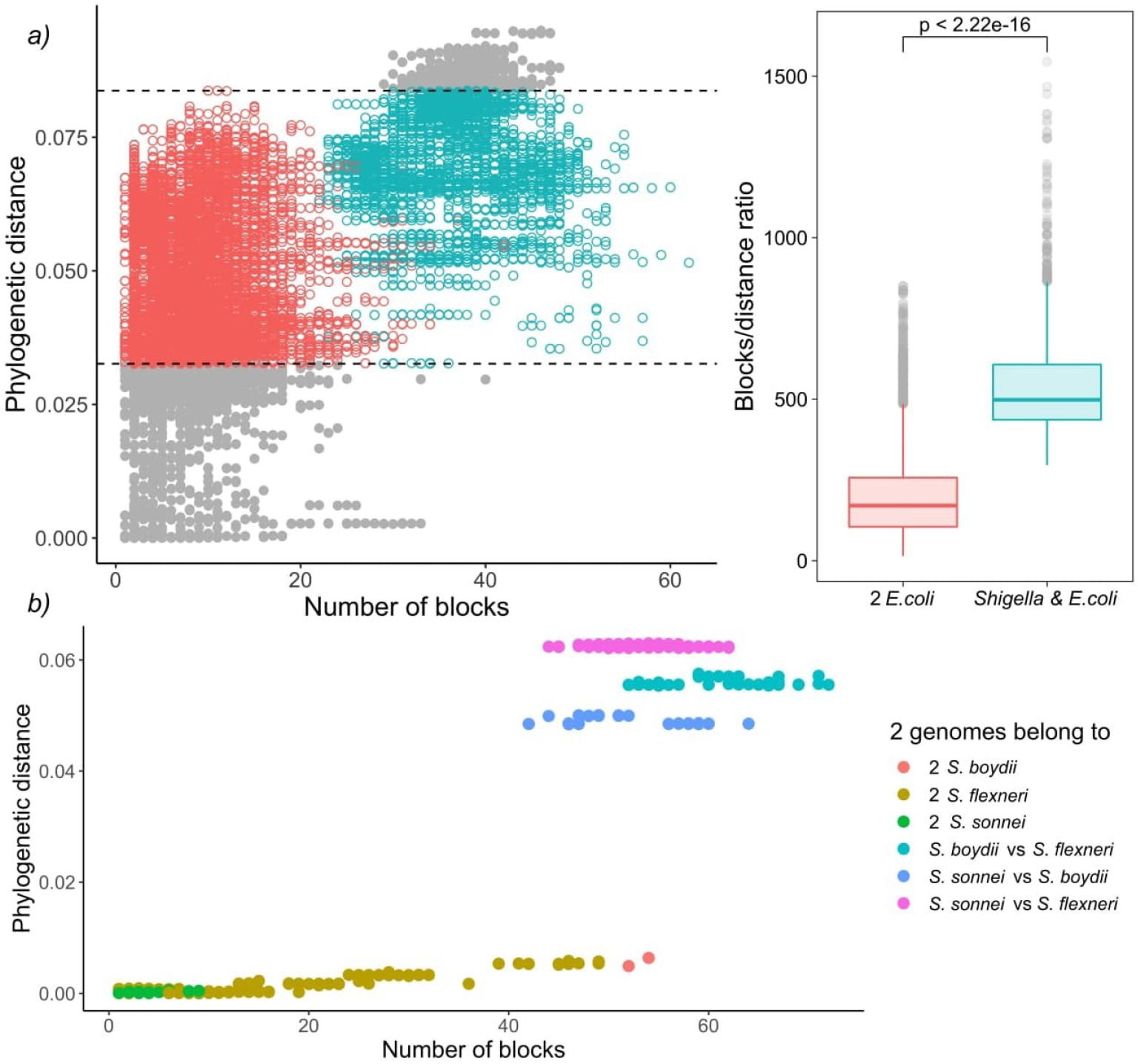

**Comparison of the ratio of phylogenetic distance to the number of synteny blocks** for two clusters from phylogroup B1. Only genome pairs in the intersection were compared. The Wilcoxon-Mann-Whitney test *p*-value is given.

### Parallel genome rearrangements

Events that occur multiple times independently on a phylogeny, called homoplasies, could indicate selection pressure acting on populations adapting to an intracellular lifestyle. Here we focus on events that have occurred several times independently in the *Shigella* lineages.

### Inversions and rearrangement hotspots

For the universal synteny blocks, we found 25 parallel inversions and 40 rearrangement hotspots where a region had been involved in inversions several times independently (Supplementary Table S3). Some of these observations were easily explained by large regions between common single-copy synteny blocks that did not allow for an accurate reconstruction of the rearrangement breakpoints (Fig. 5b). For other events, we detected the disruption of synteny by insertion sequences that had been independently integrated into the same locus and then involved in different rearrangements. For instance, independent disruption of the regions between the *pst* operon (high-affinity phosphate transport system) and the *atp* operon (proton-translocating ATPase) occurred in four independent branches and participated in four different inversions (Fig. 7). Focusing on regions that were strongly syntenic in *E. coli* but involved in rearrangements in *Shigella*, we found two adjacencies that had been disrupted independently in *S. sonnei, S. dysenteriae* and *S. flexneri*. One more interesting event is the independent inversion of the Na+/H+ antiporter gene in *S. sonnei and S. flexneri* lineages.

**Fig. 7.**
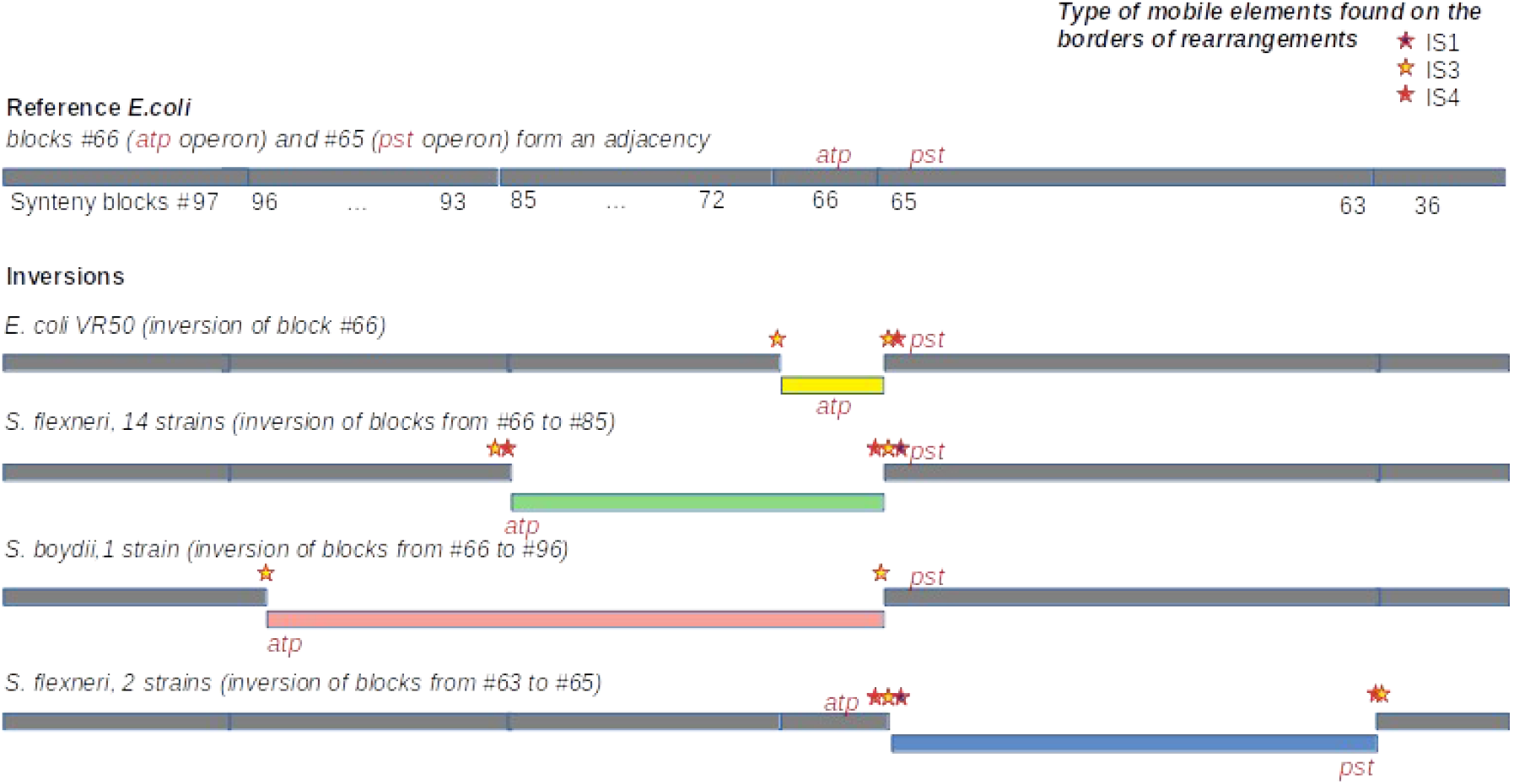

**An example of independent disruption of the adjacency** between synteny blocks #65 contained *pst* operon (high affinity phosphate transport system) and synteny blocks #66 contained *atp* operon (proton-translocating ATPase) by four different inversions. IS families found in the adjacency and putatively responsible for inversions are marked by colored stars.

We reconstructed the history of inversions in 51 strains. This dataset includes all *Shigella* spp., closely related *E. coli*, and several representative *E. coli* from each cluster without *Shigella* (Supplementary Figure S5). Of 114 reconstructed inversion events, 103 were mapped to *Shigella* branches, while only 11 corresponded to *E. coli* branches. Of these inversions, 33 were mapped to the branch separating two *S. dysenteriae*; three other *Shigella* species demonstrated rearrangements at internal and terminal branches. These results are consistent with the estimation of rearrangement rates using the number of syntenic regions as the indicator.

### Deletions

In addition to IS accumulation, *Shigella* adaptation had been accompanied by massive pseudogenization that in total resulted in genome reduction (39). These trends are well-known features of many pathogenic and symbiotic bacteria (40). Indeed, *Shigella* have significantly smaller genome size than all pathogenic and non-pathogenic *E. coli* (Supplementary Figure S6a). Note that the genome size of the two most abundant *E. coli* pathovars (ExPEC and STEC) is larger than that of non-pathogenic *E. coli* (Supplementary Figure S6b,c). Taking into account the high rate of non-homologous recombination, we anticipated seeing an increased rate of loss of non-universal synteny blocks in *Shigella*. We identified parallel insertions, deletions, and duplications in 2256 out of 3642 non-universal synteny blocks across the *E. coli / Shigella* phylogenetic tree (Supplementary Table S4). Three blocks lost in all *Shigella* and only two *E. coli* had affected the propionate catabolism (*prpABCDER*) operon. However, we did not find any strictly *Shigella-*specific large-scale deletion events.

### Insertions

In contrast, we found one single-copy synteny block that was present in *Shigella* and EIEC but absent in other *E. coli*, likely indicating acquisition of this block in *Shigella* rather than multiple independent losses in *E. coli*. This region contained the gene *IpaH1880* encoding an E3 ubiquitin-protein ligase, one of *Shigella* invasion-plasmid antigens, with a highly conserved 270 bp upstream non-coding region (Fig. 8a). In *S. sonnei, S. flexneri*, and EIEC the fragment was integrated in the same locus, while in *S. boydii* and *S. dysenteriae* this fragment was found in other loci (Supplementary Table S5). Although the *ipaH* genes are often surrounded by prophage genes and insertion sequences, they do not form stable genomic islands. Thus, the mechanism of *ipaH* integration in chromosomes is uncertain.

**Figure 8.**
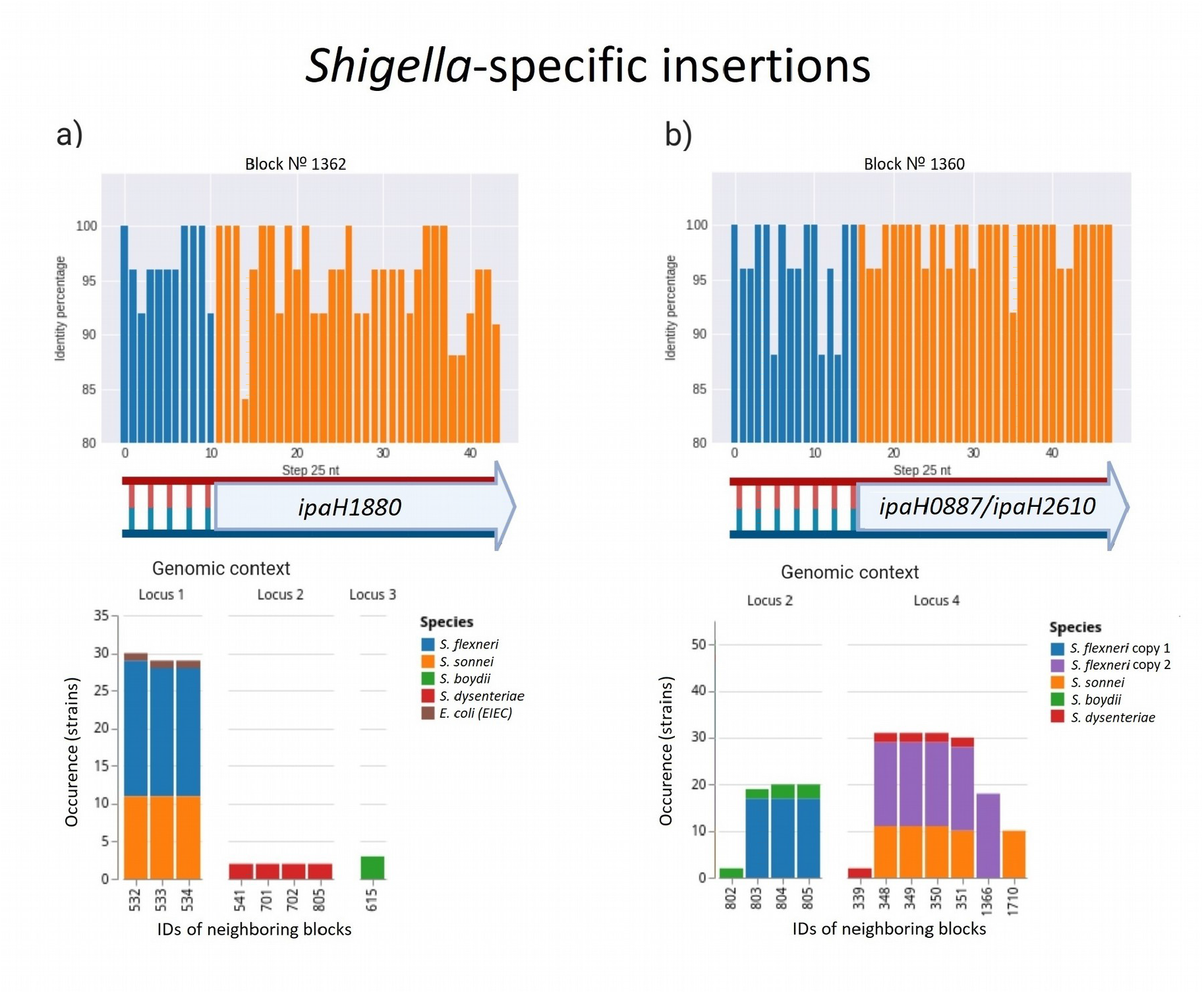
Two parallel *Shigella-specific* insertions. Both inserted blocks contain long non-coding regions with more than 85% identity and *ipaH* genes coding E-3 ubiquitin-ligases with different substrate-specific domains. Identity was calculated for nucleotide alignments of the synteny blocks separately with step 25 nt. Location of the inserted blocks in different genomic contexts confirms independent acquisition of the block by different *Shigella* lineages; in *S. dysenteriae* block #1360 was inserted twice in different loci. We took into consideration up to five of the nearest blocks, both in upstream and downstream direction, at the distance not exceeding 10 kb. Colors correspond to *Shigella* lineages. Gene composition of the neighboring blocks for both insertions is shown in Supplementary Table S5.

One more block was present only in *Shigella* but absent in other *E. coli* including EIEC. This region also contained the genes *ipaH0887/ipaH2610* encoding another E3 ubiquitin-protein ligase, again preceded by a conserved non-coding region of 230 nt (Fig. 8b). In the chromosomes of *S. flexneri* we identified two copies of this block, located at a substantial distance from each other; in some strains, one of the gene copies is marked as a pseudogene. There are two possible evolutionary scenarios explaining this block duplication. One is gene duplication in the common ancestor of *S. flexneri*; another explanation is the independent acquisition of the copies by horizontal transfer. Based on adjacent blocks, the fragments in different strains cluster into two groups; *S. sonnei* and *S. dysenteriae* have their only copy in the first locus, while *S. boydii* has its only copy in the second locus (Supplementary Table S5). This pattern may be explained by the independent transfer of the genes via site-specific insertion or via homologous recombination as these mechanisms retain the gene environment.

The evolutionary history of syntenic blocks seems relevant to the functional specificity of *Shigella* and EIEC, as the products of *IpaH* genes are secreted by intracellular bacteria via the type III secretion system (T3SS) (40) and, therefore, the *IpaH* genes repertoire may confer *Shigella-* or EIEC-specific functionality. Sequence conservation of the non-coding region indicates the importance of this sequence either for the integration or for the regulation of the *ipaH* transcription. Being highly-conserved in different *Shigella* strains, these non-coding fragments are found only in *ipaH* upstream regions and are not homologous for two different *ipaH* genes. Thus, we tentatively suggest that these non-coding fragments hold regulatory elements and play a role in *Shigella* pathogenicity.

### Rates of homologous recombination

We hypothesized that disruption of syntenic regions should decrease the rate of homologous recombination. To check this, we calculated fractions of horizontally transferred fragments in strains using pairwise genome alignments of *E. coli, S. flexneri, S. boydii*, and *S. sonnei* (see Methods). Indeed, at the same level of divergence between strains (with the Poisson λ parameter in the vertically inherited fraction ranging from 0 to 1.45), pairs of *Shigella* strains had a significantly lower fraction of fragments horizontally transferred by homologous recombination, (1–*W*)=0.094±0.017, in comparison to pairs of *E. coli* strains, (1–*W*)=0.612±0.002 (*p*=4.17×10^−124^, the Wilcoxon test) (Fig. 9).

**Figure 9.**
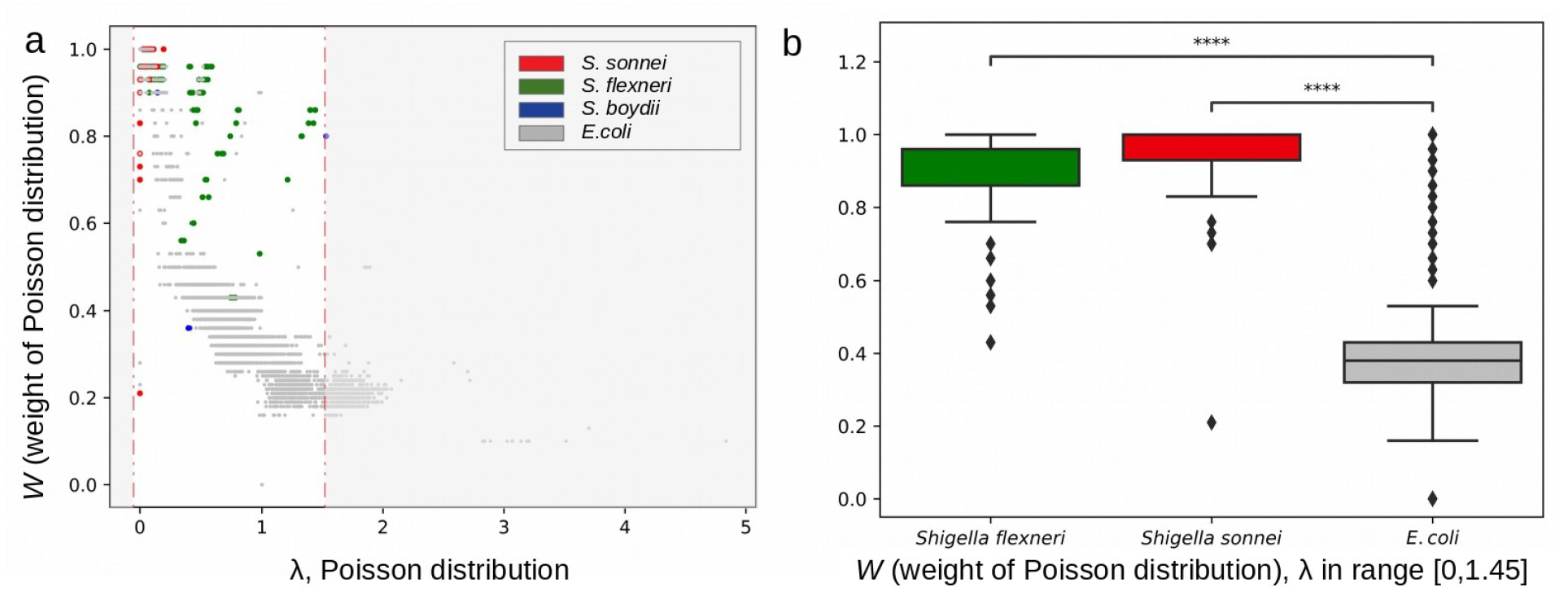
Rates of homologous recombination. **(a)** Fractions of horizontally transferred fragments in strains from the B1 phylogroup calculated as the weight of the Poisson distribution (see Methods). **(b)** The weight of Poisson distribution in *Shigella* and *E. coli* in the same interval of evolutionary distances between strains.

The estimated parameter λ may reflect many evolutionary parameters such as the generation time and the intensity of selection. However, the observation that larger Poisson lambda corresponds to lower clonal fraction of the genome, as expected, demonstrates the consistency of our results. We used this technique for *Escherichia* pairs and *Shigella* pairs at the same level of sequence similarity, and hence demonstrated that the clonal genome fraction is larger in the latter, confirming our hypothesis.

## Discussion

Compared to other pathogenic *E. coli, Shigella* (i) accumulated a large variety of ISs, (ii) acquired new chromosomal genes, (iii) experienced exceptionally high rates of intragenomic rearrangements and (iv) had a decreased rate of homologous recombination.

The diversity of *E. coli* pathotypes is explained by the high plasticity of its genome, as genes responsible for pathogenic traits are mostly acquired by extensive horizontal gene transfer and are often conveyed by mobile genetic elements (41–43). Both *Shigella* and EIEC spend much of their life cycle within eukaryotic cells and share many invasion-related functional systems. The adaptation to an intracellular lifestyle was conferred by the acquisition of the pINV plasmid encoding a type III secretion system (T3SS) (40). Phylogenetic analysis supports the hypothesis that four existing *Shigella* lineages and the EIEC strain have arisen from different ancestral *E. coli* strains on multiple independent occasions (3, 4, 7, 8).

The delivery of bacterial virulence proteins, called “effectors,” into host cells via T3SS plays a crucial role in the infection strategies of *Shigella*. Such effectors are involved in the reorganization of the host cell cytoskeleton and in the modulation of cellular signalling pathways that allow the bacteria to evade the host immune response (44). Each IpaH family protein likely has a specific host target protein due to the substrate recognition domain, and thus makes a distinct contribution to bacterial pathogenesis (40). Most T3SS effectors are encoded by plasmid *pINV* genes, while the biological role of chromosomally encoded *ipaH* genes remains obscure. Based on our results we hypothesize that some chromosomal E3 ubiquitin-protein ligases are essential for *Shigella*/EIEC pathogenicity while others may provide specific advantages. The differences in the composition of functional types of ubiquitin-protein ligases may also explain the differences in the infectious dose and disease severity between *Shigella* and EIEC pathotypes.

High numbers and the variety of mobile elements in evolutionarily young intracellular pathogens is usually explained by weaker selection against repetitive elements due to decreased effective population size (9, 17, 38). IS elements may drive the evolution of chromosome organisation by causing non-homologous recombination (45). Genome rearrangements change the chromosomal architecture, which can alter gene composition and gene expression. These events are largely detrimental for free-living bacteria and are expected to be selected against (18). On the other hand, in a new environment, non-homologous recombination and the associated functional changes may provide a base for adaptation (46, 47). For instance, *Burkholderia mallei*, a young obligate intracellular pathogen, has lost numerous clusters of genes through IS-mediated elimination as demonstrated by the comparison of its genome to strains of the ancestral species, *Burkholderia pseudomallei* (48, 49). The genome reduction of *B. mallei is* up to 30% in some strains and the adaptation is still not complete (50). In comparison to other *E. coli, Shigella* have slightly smaller genomes and the evolution was accompanied by IS-mediated pseudogenization, but not large-scale deletions.

IS families differ in the expansion rate in *Shigella* lineages, which is expected given the independent origin of these lineages. Two types of IS (IS3, IS4) revealed high expansion rates in all *Shigella* lineages; this observation is in agreement with a recently published analysis of draft *Shigella* and EIEC genomes (13). These ISs are not common for *E. coli* populations but are a part of *pINV* that explains their expansion after the plasmid acquisition. In contrast, IS1 is present in many pathogenic *E. coli* and the difference in its frequency in pathogenic *E. coli* and *S. sonnei* is not statistically significant. The number of IS elements and, consequently, the rate of genome rearrangements in the EIEC strain were comparable with those in other *E. coli*. On the other hand, draft EIEC genomes featured larger IS frequencies in EIEC populations in comparison to other *E. coli*, but lower than in *Shigella* strains (13); but the rates of genomic rearrangements were not estimated in the cited paper as the studied genomes had not been assembled. Taken together, we propose that differences in the number of IS genomic elements may have influenced different stages of formation of intracellular pathogens EIEC and *Shigella* spp.

An expected consequence of frequent genome rearrangement is a decrease in the rate of homologous recombination. Indeed, in comparison to *E. coli, Shigella* genomes contain fewer DNA segments horizontally transferred by homologous recombination. However, this also could be explained by smaller population size and isolated intracellular lifestyle of *Shigella* strains. Homologous recombination levels in core genes, manifesting as incongruence of gene phylogenetic trees with the strain phylogeny, is smallest in endosymbionts and intracellular pathogens (51). These are likely interconnected processes. Bottlenecks and decreased selection pressure lead to the increase in the number of IS elements (9, 50); this in turn provides more opportunities for genome rearrangements that become tolerated due to decreased selection. Indeed, bursts of rearrangements were observed in the genomes of pathogens that had recently changed host and lifestyle, such as *Yersinia pestis* (48) and *Burkholderia mallei* (52). On the other hand, relative isolation of strains with the mainly intracellular lifestyle provides fewer opportunities for homologous recombination, while the lack of genome collinearity creates mechanistic obstacles to the process (53). At that, neither increased rearrangement rate, nor decreased homologous recombination rate are observed in intracellular EIEC strains of *E. coli*, supporting the link between these phenomena.

The analysis of genome rearrangements requires complete genomes, while less than 1% of available *Shigella* genomes have sufficient quality of assembly. Another issue is that misassemblies are mainly caused by genomic repeats and may be confused with true rearrangements. Although there are several strategies widely used for assembly validation such as long read (re)sequencing and/or PCR contiguity verification (54–56), some of the detected rearrangements may be caused by inaccuracies in gap closure procedures. While each particular observation requires experimental verification, the observed correlation between rearrangement rates and mutation rates in closely related strains allow us to conclude that assembly errors do not affect the evolutionary signal in the analysed data (52).

Due to high genomic plasticity, pathogenic *E. coli* are among the most frequent causes of bacterial infections in humans (57, 58). In particular, the *E. coli* O104:H4 outbreak in Germany in 2011 was caused by a strain that had acquired characteristics of two previously described pathotypes (59). The emergence of antibiotic-resistant *Shigella* with the absence of an effective vaccine highlights the importance of a detailed investigation of this pathogen (60). Our results contribute to the understanding of genomic properties associated with adаptation to intracellularptation to intracellular lifestyle of *Shigella* and EIEC and the developed approaches should be broadly applicable to other young bacterial pathogens.

## Author statements

### Authors and contributors

MSG and OOB conceived the study, ZS, OOB, and MSG designed the study; ZS, RA, AZ, and NA developed the methods, ZS, YY, RA, ND and OOB analyzed the data; ZS, NA, OOB, and MSG wrote the manuscript. All authors read and approved the final version of the manuscript.

### Data summary

The datasets supporting the conclusions of this article and used scripts are available at https://github.com/zseferbekova/ShigellaProject. The authors confirm all supporting data have been provided in the article or as supplementary data files.

### Ethical approval

Not applicable

### Consent for publication

Not applicable

## Acknowledgements

We thank Fyodor Kondrashov for valuable advice and manuscript proofreading. We also thank Alla Mikheenko for assistance with Circos.

---

**Figure S1. Phylogenetic trees. (a)** *E. coli* phylogenetic tree. The tree is based on the nucleotide alignment of universal single-copy orthologs. Strains shown in red were excluded from further analysis. **(b)** *E. coli/Shigella* phylogenetic tree. The tree is based on the nucleotide alignment of universal single-copy orthologs with 95% identity threshold.

**Figure S2. Construction of breakpoint graph. (a)** Genome graphs of unichromosomal circular genomes *P = (0, 1, 2, 3, 4, 5)* and *Q = (0, 1, -4, -3, -2, 5)*, the adjacency edges of the genome *P* (left) are shown in blue, the edges of the genome *G* (right) are shown in red **(b)** The breakpoint graph *G(P,Q)* of genomes *P* and *Q* **(c)** The multiple breakpoint graph of five unichromosomal circular genomes.

**Figure S3. Chromosome maps (a)** *Escherichia coli* C4 **(b)** *Escherichia coli* cfsan029787 **(c)** *Shigella flexneri* 1a 228 **(d)** *Shigella boydii* atcc 9210 **(e)** *Shigella dysenteriae* sd197 **(f)** *Shigella sonnei* fdaargos 524. The inner circle — GC-skew, the second (blue) circle — synteny blocks, the third (green) circle — universal synteny blocks, the fourth (grey) circle — ISs, the fifth (grey) circle — density of ISs.

**Figure S4. Phylogenetic distance versus the number of synteny blocks** for each pair of genomes **(a)** in phylogroup A, **(b)** in phylogroup B2, **(c)** in phylogroup D, **(d)** in phylogroup F. Each point represents a pair of genomes and is coloured according to the genomes the pair includes.

**Figure S5. Inversion events reconstructed by MGRA software. (a)** A cladogram with numbers of inversions shown for each branch. **(b)** The corresponding phylogenetic tree. Phylogroups are marked with coloured strips, pathogenic strains are shown in different colours. Both trees are unrooted. The trees are also available online (iTOL): https://itol.embl.de/tree/9318063252480721583313649.

**Figure S6. Genome sizes** in **(*a*)** *Shigella*, non-pathogenic *E. coli*, other pathogenic *E. coli* and all *E. coli* (excluding *Shigella* spp); **(b)** two most abundant pathovars — ExPEC and STEC, non-pathogenic *E. coli* and all *E. coli* (excluding *Shigella* spp). **(c)** different groups of *E. coli*. The numbers above boxplots indicate the number of genomes in each group. The Wilcoxon-Mann-Whitney test with the Bonferroni correction: (*) *p* ≤ 0.05, (**) *p* ≤ 0.01; (***) *p* ≤ 0.001; (****) *p* ≤ 0.0001.

**Table S1. The list of *E. coli* and *Shigella* genomes included in the analyses. Table S2. Coordinates of synteny blocks**.

**Table S3. Adjacencies of common synteny blocks. Table S4. Copy number of non-common synteny blocks**.

**Table S5. Gene composition of synteny blocks surrounding the *Shigella*-specific insertions**.

