## Supplementary figures and images for "High rates of genome rearrangements and pathogenicity of *Shigella* spp"

### Supplementary Figure S1a

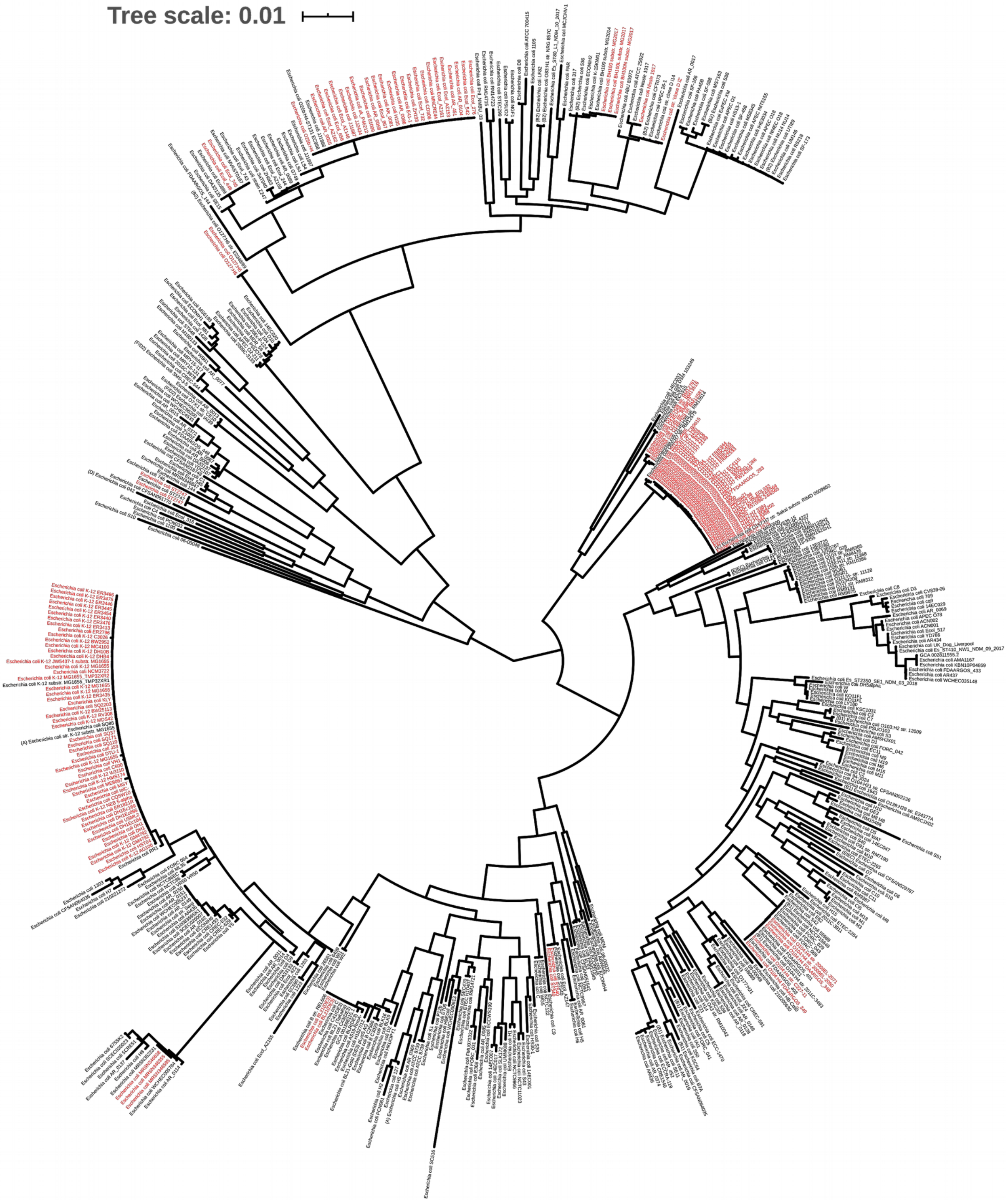

### Supplementary Figure S1b

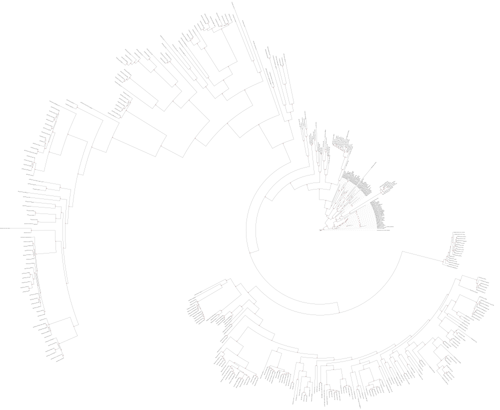

### Supplementary Figure S2

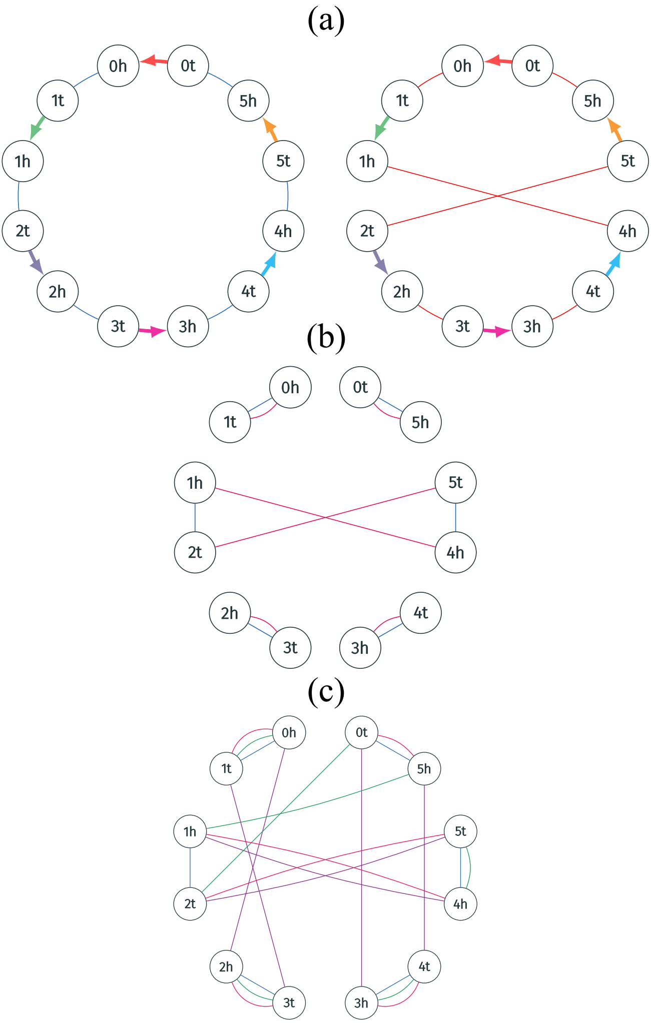

### Supplementary Figure S3

(a) Escherichia coli C4

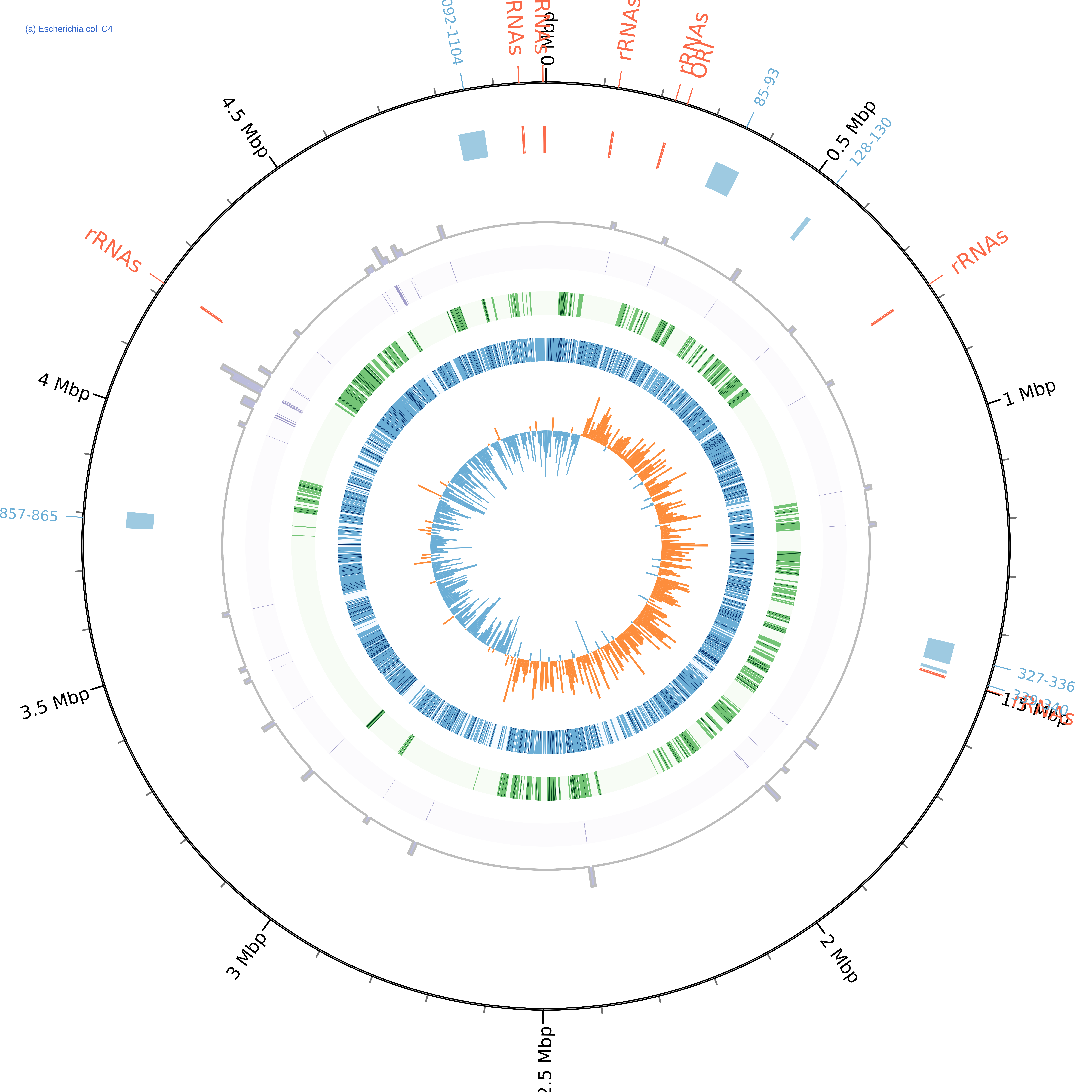

b) Escherichia coli cfsan029787

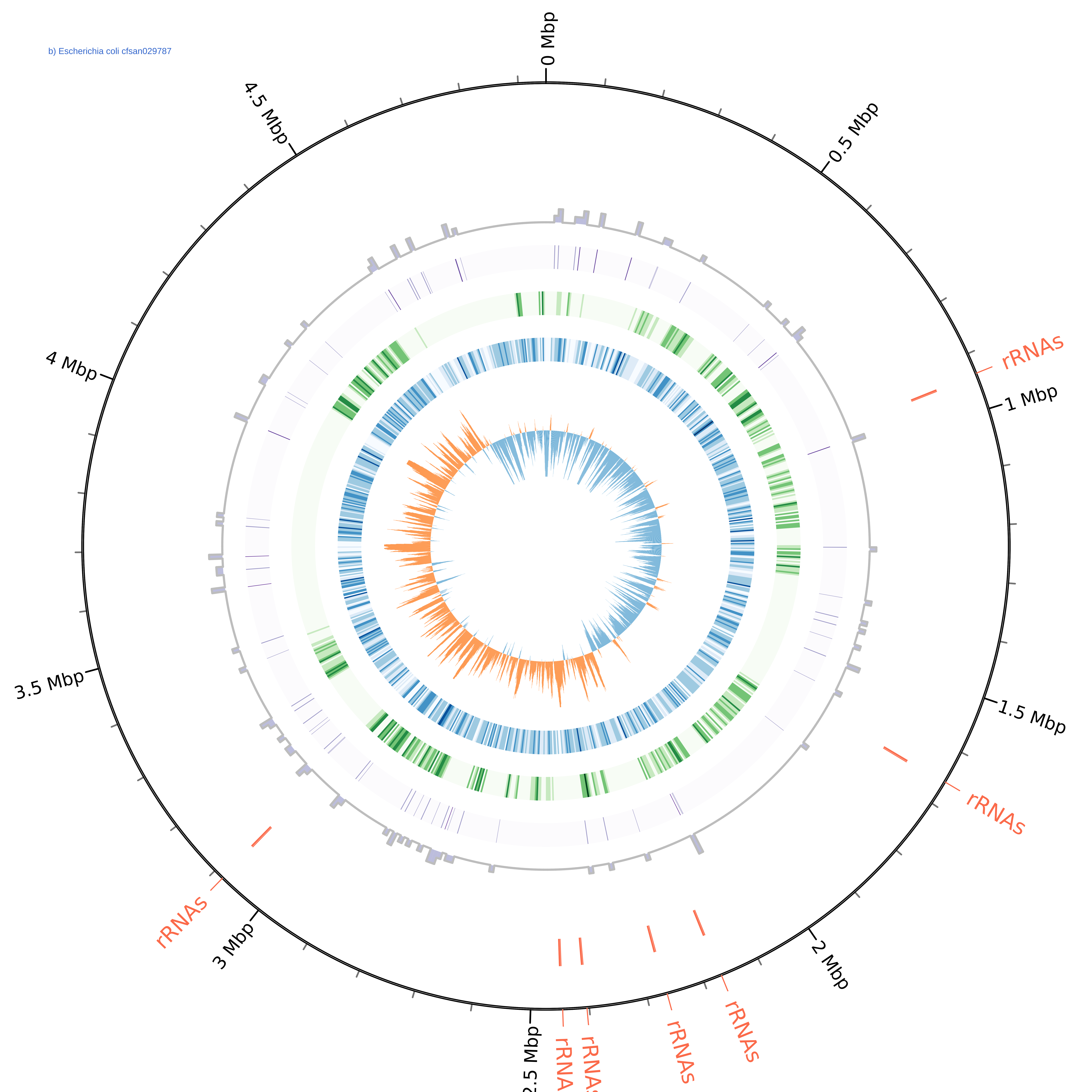

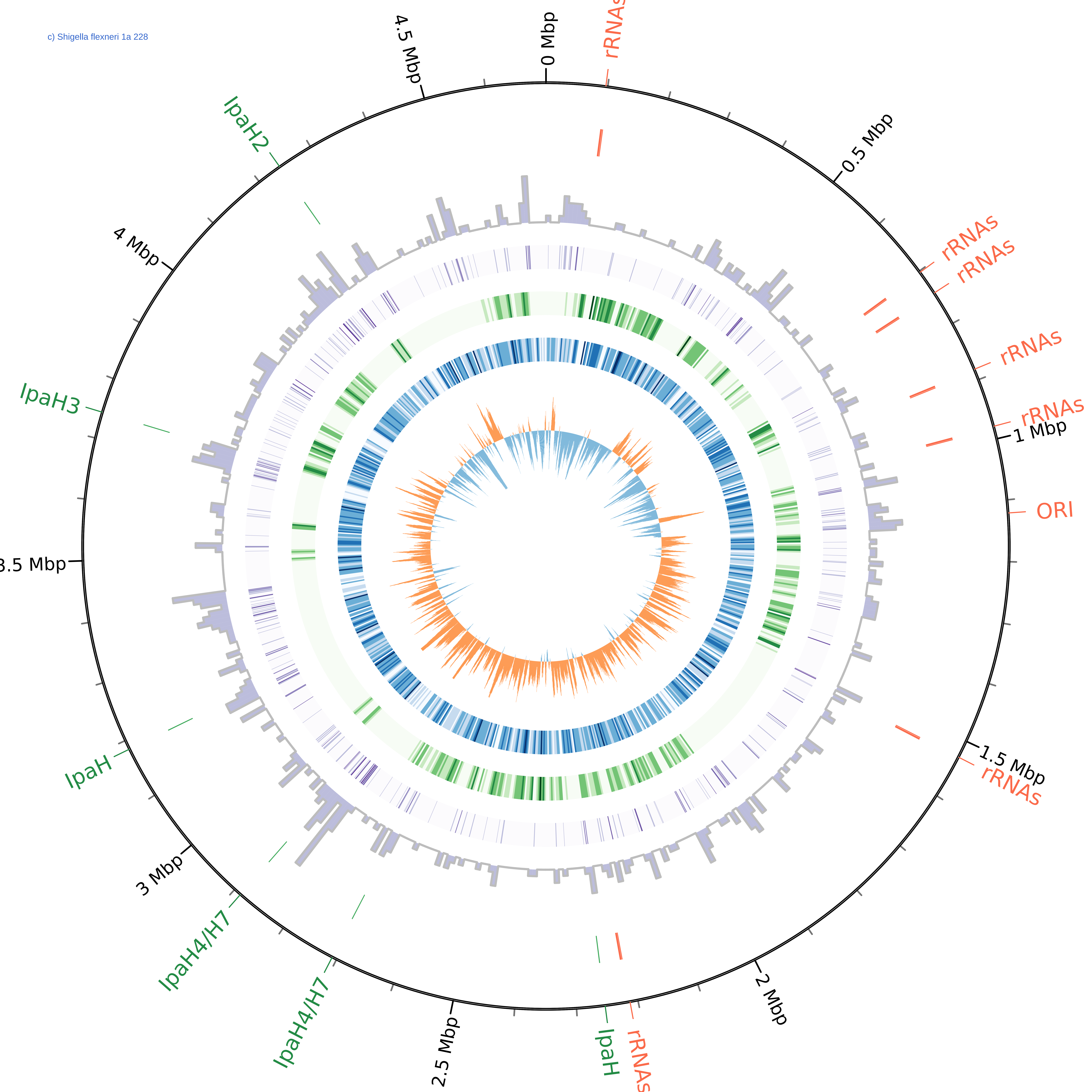

d) Shigella boydii atcc 9210

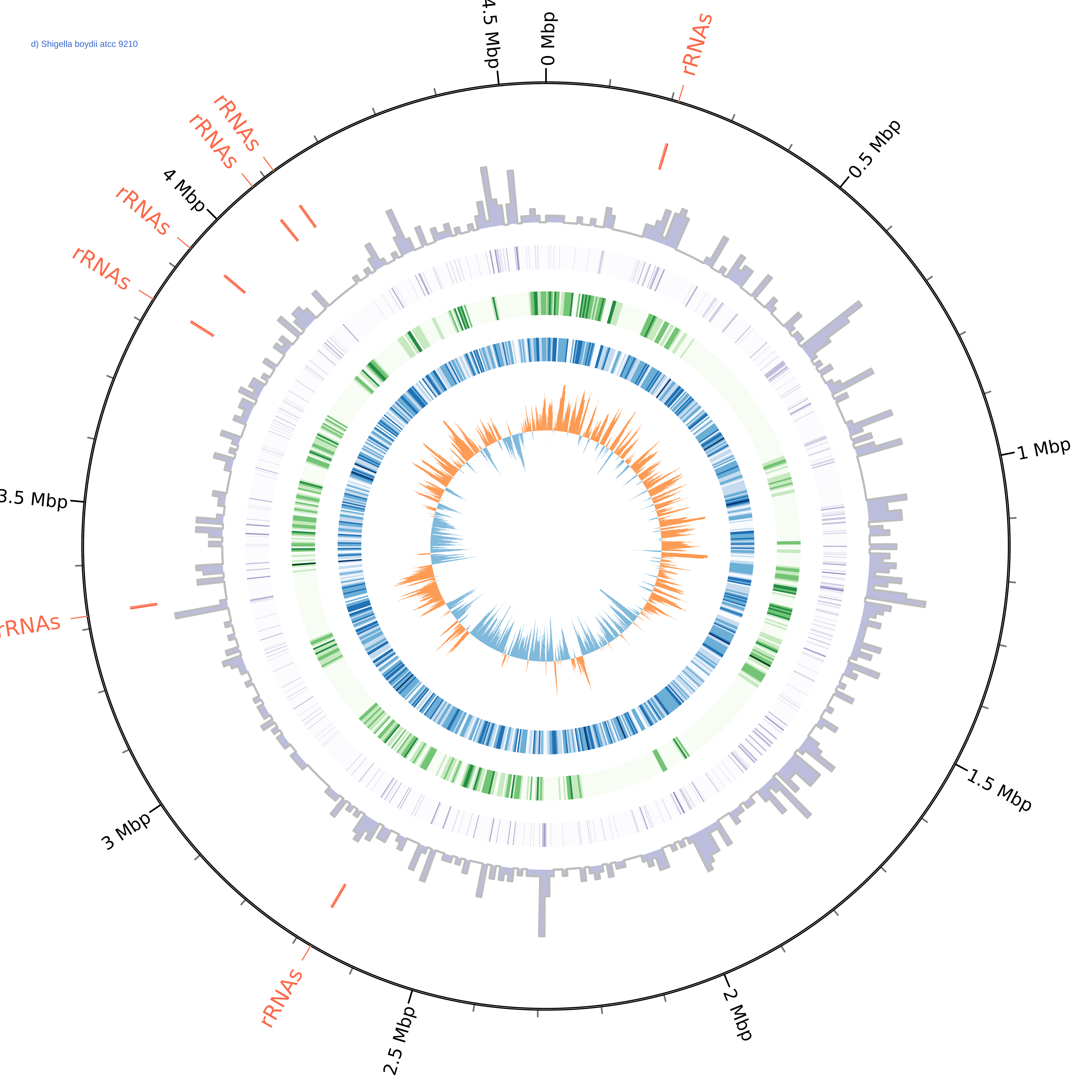

e) Shigella dysenteriae sd197

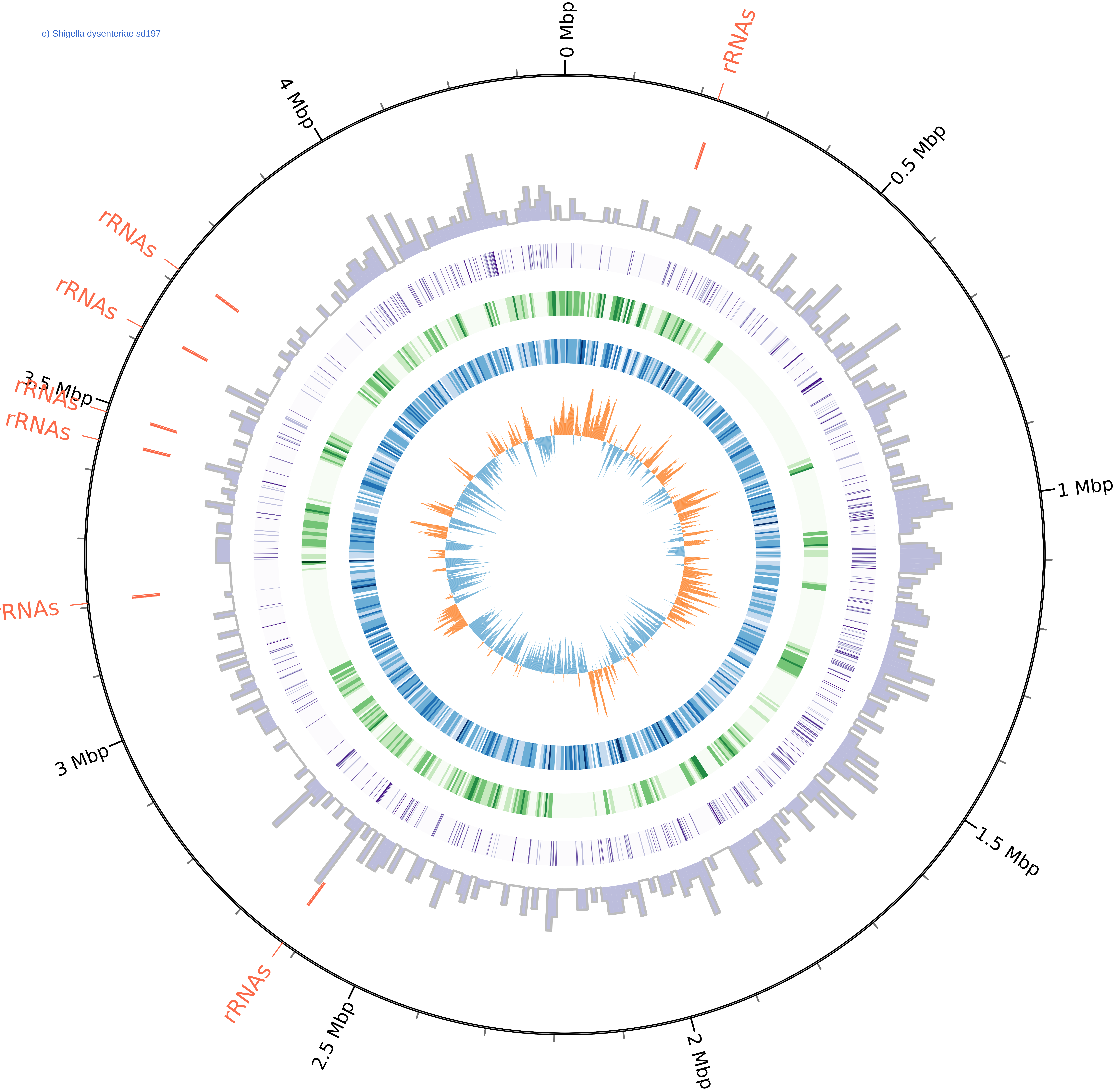

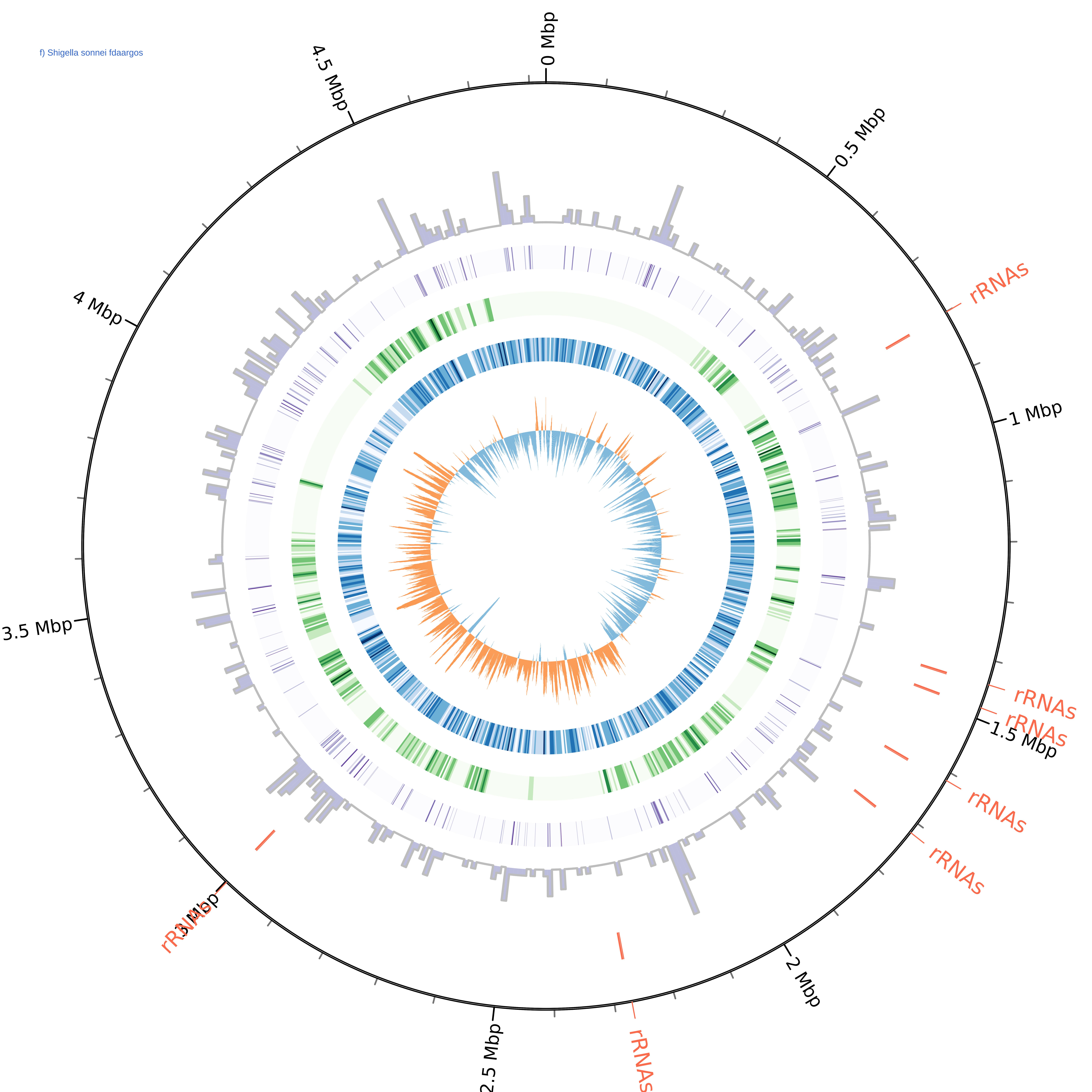

### Supplementary Figure S4

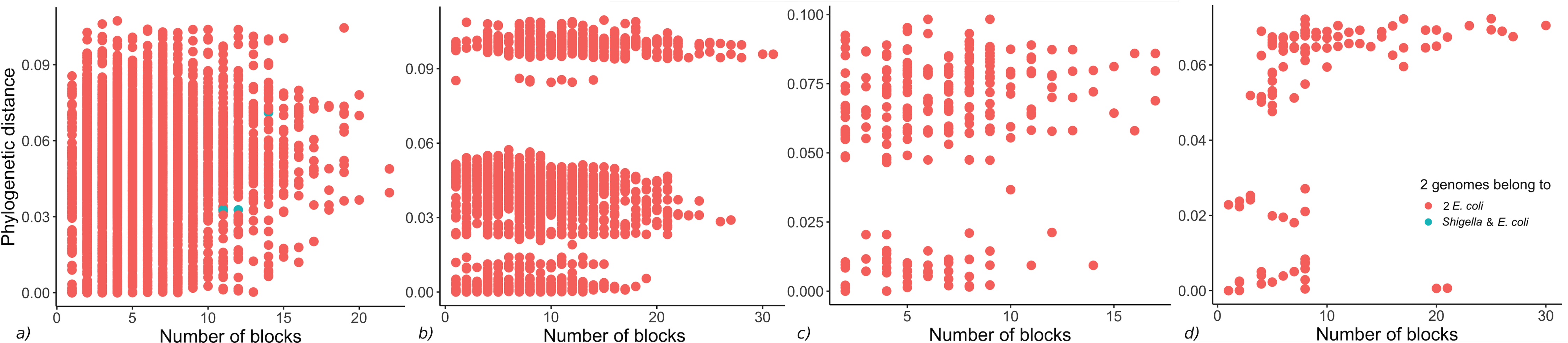

### Supplementary Figure S5

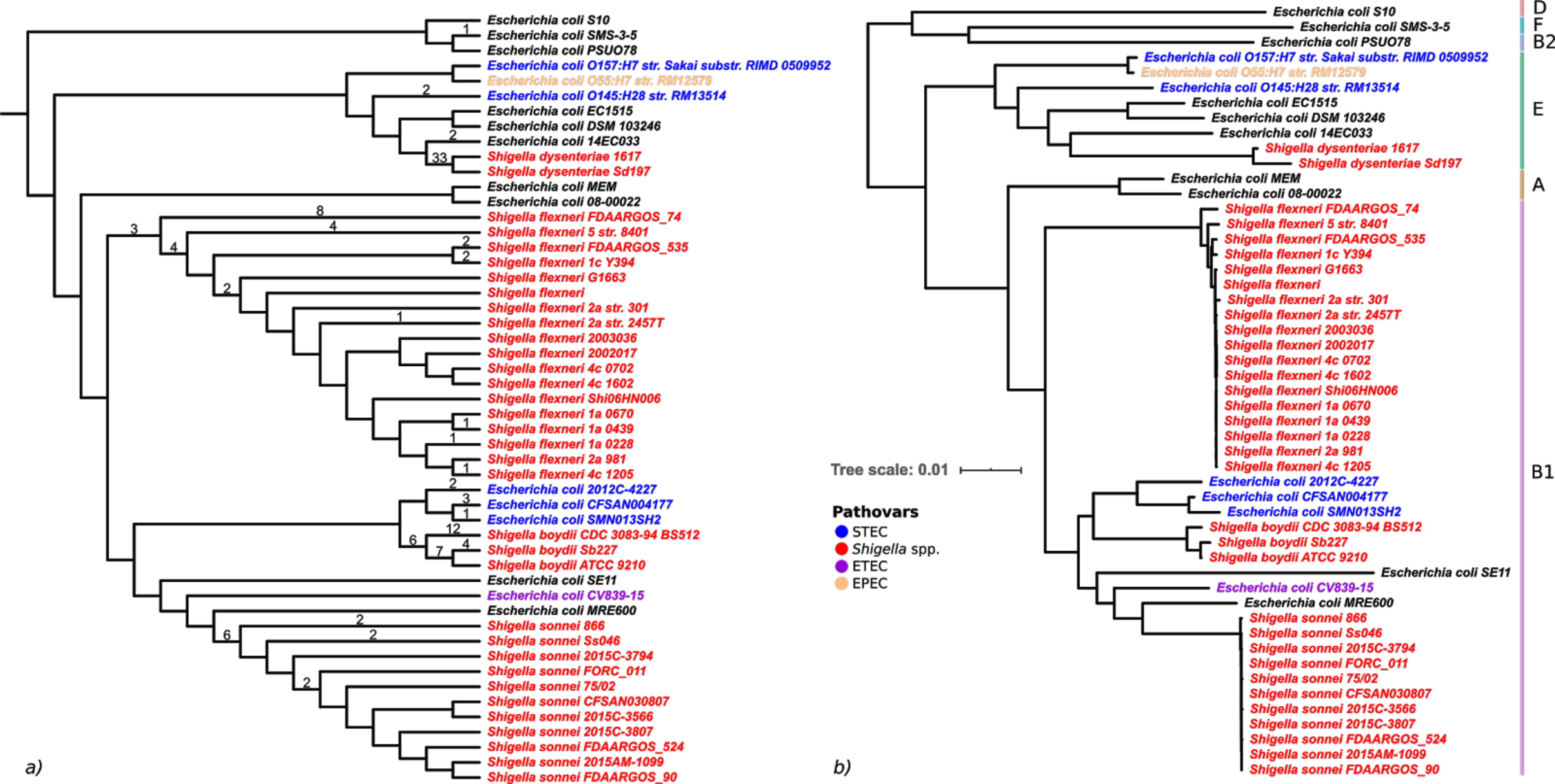

### Supplementary Figure S6

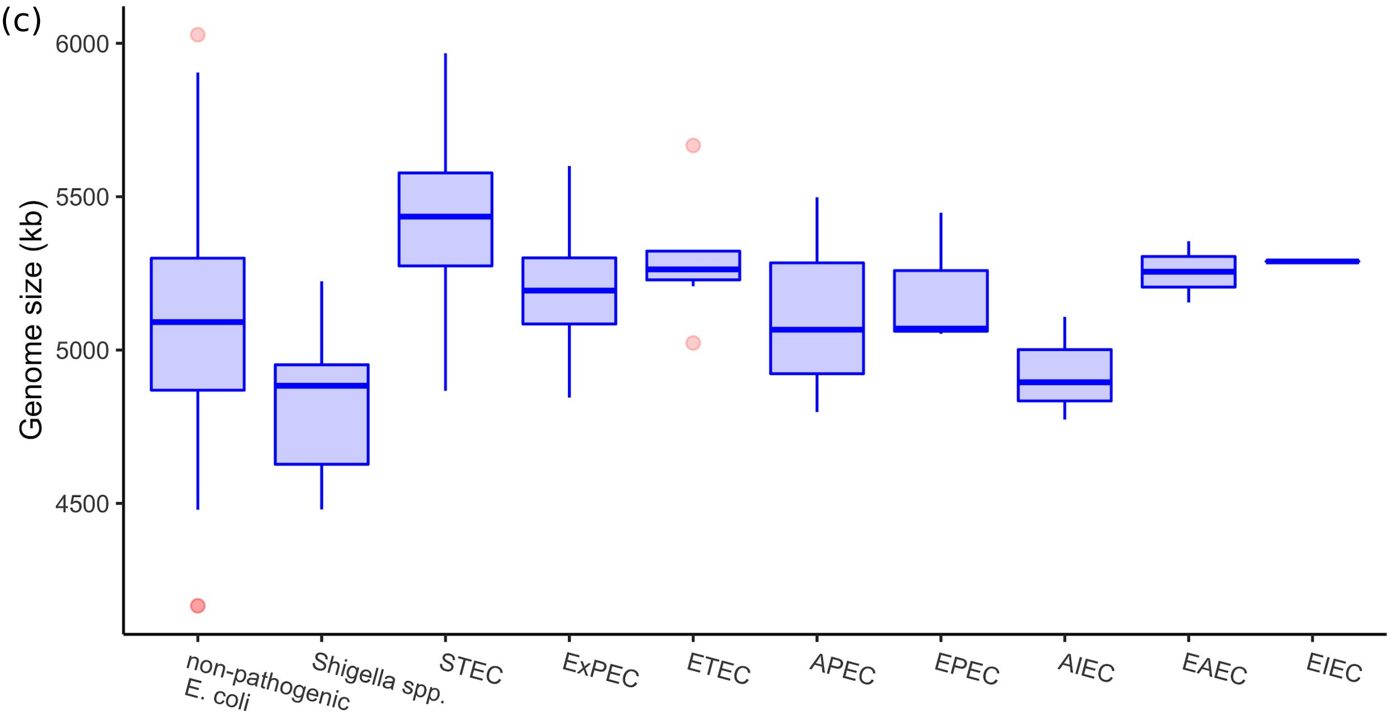
